# Compromised striatal structure and function in mouse models of RARB-related disorder

**DOI:** 10.64898/2026.02.20.706764

**Authors:** Nicolas Zinter, Devanshi Shah, Victorine Artot, Nicolas Lemmetti, Hanna Semaan, Rodolphe Soret, Nicolas Pilon, Christina Nassif, Marie-Christine Birling, Amrita Raja-Ravi-Shankar, Valérie Fraulob, Véronique Caron, André Tremblay, Gerardo Zapata, Marc Danik, Jacques L. Michaud, Wojciech Krezel

## Abstract

Dominant variants in the *retinoic acid receptor beta* (RARB) gene cause a complex disorder known as RARB-related disorder (RARB-RD), characterized by multiple congenital anomalies, global developmental delay, and dystonia. RARB-RD variants have been classified as either gain-of-function (GOF) or dominant-negative (DN) based on their cell-based transcriptional responses to retinoids. To investigate the mechanisms underlying this disorder, we generated mouse models carrying either the p.R387C or p.L402P RARB-RD variant, previously categorized as GOF and DN, respectively.

Homozygous mice for either RARB-RD variant died perinatally with colonic aganglionosis, while heterozygous mice survived and recapitulated several features of RARB-RD. In addition to microphthalmia, both *Rarb^R387C/+^*and *Rarb^L402P/+^* mice exhibited progressive coordination deficits, increased active-phase locomotor activity, and cognitive impairment in the novel object recognition test. In contrast, mice heterozygous for a null allele of *Rarb* (*Rarb^+/-^*) did not display these abnormalities. In the brain, *Rarb* is predominantly expressed in the two major populations of projection neurons of the striatum recognizable by the expression of dopamine receptors D1R/*Drd1* and D2R/*Drd2*. Marker analysis revealed a reduction in *Drd2*-expressing neurons without changes in *Drd1*-expressing neurons in both RARB-RD models. Furthermore, RARB-RD mice showed partial resistance to the cataleptic effects of haloperidol, a D2R-specific antagonist. These behavioral, cellular, and dopaminergic deficits—though not the cognitive impairments—have previously been observed in *Rarb^-/-^* mice.

To determine whether the *in vitro* effects of RARB-RD variants correlate with distinct transcriptional signatures *in vivo*, we compared the striatal transcriptome of *Rarb^R387C/+^*, *Rarb^L402P/+^*, *Rarb^-/-^* and *Rarb^+/-^*mice with their littermate controls. We found that the heterozygous RARB-RD variants and the homozygous null allele affected a large subset of common genes, with putative direct RARB targets predominantly downregulated. Notably, the transcriptional impact of the RARB-RD variants was more profound than that of the null allele, regardless of zygosity. Additionally, transcriptional changes in RARB-RD mice extensively overlapped with those observed in mouse models of Huntington’s disease, suggesting shared mechanisms affecting neuronal survival in the striatum.

We conclude that the p.R387C and p.L402P variants similarly compromise striatal integrity and function, likely through a DN mechanism. Progressive emergence of most neurologic deficits highlights a potential therapeutic window. Our results support the development of strategies aimed at silencing RARB-RD alleles.

## Introduction

All-*trans*-retinoic acid (ATRA), the active form of vitamin A, is a small molecule required for normal embryonic development^1–4^ and tissue homeostasis, especially in the brain where it regulates multiple processes^5–9^. Both loss^10,11^ and gain^12–15^ of ATRA signaling are toxic, highlighting the importance of its tight regulation. ATRA signaling is mediated by retinoic acid receptors (RARA, RARB, RARG), which act as ATRA-dependent transcription factors by heterodimerizing with retinoid-X-receptors (RXRA, RXRB, RXRG). All RARs share a common modular structure with four functional domains, including highly evolutionary conserved DNA-binding^16,17^ and ligand-binding domains^18^.

We have described a RARB-related disorder (RARB-RD, also known as MCOPS12; OMIM #615524) that is characterized by developmental eye anomalies, other congenital defects, global developmental delay, and severe motor impairment associated with dystonia^19–21^. The vast majority of RARB-RD variants are missenses clustered within the ligand-binding domain. These variants can be classified as gain-of-function (GOF) and dominant-negative (DN) variants, depending on whether their transcriptional response to 1µM ATRA in cultured cells was significantly higher or lower than that of wild-type RARB. DN variants reduce protein activity possibly by altering ligand affinity or disrupting interactions with coactivators. It remains unknown whether RARB-RD variants produce the same effects *in vivo*.

In the mouse and human brain, *Rarb* is predominantly expressed in the striatum, a highly conserved forebrain structure which receives neuromodulatory inputs from many brain regions to coordinate motor and cognitive behaviors. In humans, impaired striatal function causes several clinical manifestations, including cognitive deficits and movement disorders such as dystonia. *Rarb* is expressed in the two main neuronal populations of the striatum, namely the striatonigral (snMSNs) and striatopallidal (spMSNs) medium spiny neurons, which are recognizable by the expression of dopamine D1 receptor (D1R/*Drd1*) and dopamine D2 receptor (D2R/*Drd2*), respectively^22,23^. *Rarb* is continuously expressed in these neurons from their birth. Mice homozygous for a null allele of *Rarb* (*Rarb^-/-^*) display a distinct pattern of motor coordination deficits that are consistent with striatal dysfunction^24^. Loss of *Rarb* function disrupts snMSNs proliferation and differentiation during embryogenesis^2^ whereas it leads to a reduction in the number of spMSNs beginning postnatally^22^. Decrease in *Rarb* expression induced in the adult striatum also leads to coordination deficits and a loss of spMSNs, further supporting a postnatal role of RARB in maintaining these cells^22^. All together, these observations strongly suggest that the neurodevelopmental features of RARB-RD are mainly explained by the disruption of RARB function in striatal neurons.

To further understand the molecular and cellular mechanisms underlying RARB-RD, we generated mouse models carrying the pathogenic variants p.(Arg387Cys) (p.R387C) and p.(Leu402Pro) (p.L402P). p.R387C is found in approximately 30% of all known individuals with RARB-RD and p.L402P was reported in multiple unrelated individuals with this disorder^19^. Both Arg387 and Leu402 are located in the ligand-binding domain; notably, p.L402 is also part of helix 12, which plays a direct role in co-activator recruitment. Functionally, p.R387C and p.L402P exert GOF and DN transcriptional effects, respectively, in cultured cells supplemented with 1µM ATRA^19–21^. Here, we show that, despite their structural and functional differences, these variants similarly affect behavior as well as dopamine signaling and transcriptional regulation in the striatum. This convergence implies that they may disrupt RARB signaling through a shared DN mechanism. Finally, we found that the transcriptional signatures associated with RARB-RD variants share remarkable similarities with those observed in Huntington’s disease (HD), suggesting the existence of common mechanisms underlying these two disorders.

## Materials and Methods

### Animals

Mice carrying the NM_011243.2:c.1159C>T (NC_000080.7:6037504C>T; GRCm39/mm39) and NM_011243.2:c.1205T>C (NC_000080.7:g.6037550T>C; GRCm39/mm39) variants, coding for the RARB protein variants p.R387C and p.L402P, respectively, were generated using CrispR/Cas9 at the McGill Integrated Core for Animal Modeling (p.R387C) and the Institut Clinique de la Souris (PHENOMIN-ICS; IGBMC) (p.L402P). Briefly, gRNAs, designed via *Crispor*^25^, were complexed with the Cas9 protein and microinjected into C57Bl/6 Envigo embryos along with a single-stranded oligodeoxynucleotide carrying the respective variant. The microinjected embryos were then transferred into CD-pseudo-pregnant females, resulting in live pups. The pups were screened via Sanger sequencing, and F0 mice with the desired variant were identified (Fig. S1A,B). The lines were backcrossed over six generations on the C57BL/6NCrl backgrounds. Genotyping of the variants was performed as described in Fig. S1C. *Rarb^+/-^* mice have been previously described ^22,26^.

All mice were housed under 12h/12h light/dark cycles with the beginning of the light period at 7am and had access to food and water *ad libitum*. Experiments were approved by the institutional animal care committees of CHU Sainte-Justine Azrieli Research Center and IGBMC as well as by the French Ministry for Superior Education and Research (projects APAFIS #35396-2022021019043817 and APAFIS #39604-2022103116422358).

### Behavioral phenotyping

Both male and female mice were tested using a battery of behavioral tests according to standard operating procedures described in^22,27^. Tests were carried out in the following order: open-field test, rotarod, spontaneous locomotion in actimetric cages, grip test, hanging test, notched bar, elevated plus maze, haloperidol induced catalepsy. In absence of any significant differences between sexes, data from males and females were pooled for analyses (detailed methods in the supplementary material).

### Pharmacology

Experiments involving haloperidol (Sigma-Aldrich, France) administration followed by cFOS analyses were performed as previously described^22^. Pramipexol (CliniScience, France), dissolved in DMSO and saline, was administrated at doses of 0.2 or 2 mg/kg via intraperitoneal injection 30 minutes prior to a 10 minute-long open-field assay, followed by 3 trials of the rotarod assay. For spontaneous locomotor activity analysis, mice were placed in actimetric cages for a habituation period (from 11:00 a.m. to 7:00 p.m.), injected with pramipexol at 7:00 p.m. and then returned to the actimetric cages for 24 hours with free access to food and water.

### In situ hybridization and cell counts

Rostral cryosections, 14 µm thick (at Bregma +1.34 mm), were processed and used for *in situ* hybridization and cell counting, as described^22^.

### Western blotting

Striatum samples from newborn (P0) and adult (20 weeks old) mice were prepared and run as described^28^. For antibodies and their dilutions, see the supplementary material section.

### Immunofluorescent detection and cell quantification

Immunofluorescent detection of RARB and cFOS on brain sections (Bregma +1.34mm – 1mm) was performed as described in^22,28^. Immunofluorescent detection of NEUN, GFP, and RFP on corresponding regions of floating brain sections (collected from perfused brain samples) were performed as described^29^. All preparations were imaged using a Leica TCS SP8 confocal microscope. Antibodies are listed in the supplementary material section. The number of cells labelled by the different antibodies was counted without prior knowledge of genotype and gender. The distribution of all neurons (NeuN^+^), neurons expressing the D1 receptor (RFP^+^), and neurons expressing the D2 receptor (GFP^+^) was analyzed automatically using the ImageJ software. The cell counts from each animal (4 animals/group) correspond to the average of measurements obtained from three adjacent slices.

### Transcriptomic studies

The dorsal striatum was dissected from 1 mm-thick coronal slices obtained from the brain of P60 male mice. Total RNA was purified from samples using the RNeasy Mini Kit with on-column DNase digestion (Qiagen). Ribosomal RNA was depleted from each mouse total RNA sample (250 ng) (NEBNext rRNA depletion kit, New England BioLabs) for the preparation of rRNA-depleted (HMR) stranded libraries (Illumina). Paired-end (2 x 100) sequencing was performed on an Illumina NovaSeq 6000 System (Illumina) with number of reads per sample ranging from 52 to 89M (average = 72M). Bioinformatic processing of the raw data was performed using the GenPipes framework by the Canadian Centre for Computational Genomics (McGill University, Montreal)^30^. We confirmed the genotype of each sample by searching for the presence of the mutation of interest in the *Rarb* transcripts. The transcriptome of mice with a given mutation was compared to that of littermate wild-types (N for each group = 4-5). To define a gene as being differentially expressed, we utilized a FDR-corrected Q-value cutoff of 0.05. Gene overlap analysis was performed with GeneOverlap^31^ gene set enrichment analysis with GSEA^32^ and pathway enrichment analysis with Enrichr^33^.

### Statistical analyses

Pairwise comparisons were analyzed using the Mann-Whitney test as the data did not follow a normal distribution according to the Shapiro-Wilk test. The comparisons of *Rarb^+/+^*, *Rarb^R387C/+^* and *Rarb^L402P/+^* adult mice were carried out using one-way ANOVA with genotypes as a factor. When measures were repeated on the same subjects (distance and rearing activity in openfield, locomotor activity in actimetric cages, longitudinal analyses), analyses were performed using repeated-measures ANOVA. Pharmacological treatments analyses were carried out using two-way ANOVA with treatment and genotypes as independent variables. All post-hoc analyses were performed using Bonferroni’s multiple comparisons test.

## Results

### Mouse models of RARB-RD recapitulate its key manifestations

To investigate the mechanisms underlying the development of RARB-RD, we generated mice carrying the p.R387C and p.L402P pathogenic variants. We found that, in contrast to *Rarb^-/-^*mice, which survive into adulthood, mice homozygous for either mutation died early in the neonatal period. In both cases, autopsies revealed a dilation of the mid-colon combined with narrowing of the most distal segment of the colon - a phenotype that has not been reported in *Rarb^-/-^* mice. Characterization of the enteric nervous system (ENS) in *Rarb^R387C/R387C^* newborns using βIII-tubulin immunofluorescence revealed the absence of the characteristic lattice-like network of neural ganglia observed in littermate controls (Fig. S2A). Moreover, colonization of the colon by βIII-tubulin+ neural-crest derived ENS progenitors was significantly less extensive in mutant embryos than in littermate controls from E12.5 onward (Fig. S2B). These anomalies are typical of the a-/hypo-ganglionosis found in mouse models of Hirschsprung disease. Interestingly, enhancer variants in the *RET* gene disrupting RARB binding have been shown to increase the risk of Hirschsprung disease^34,35^. Consistent with this observation, we observed fewer RET-expressing cells in *Rarb^R387C/R387C^*embryos compared to controls (Fig. S2C). Taken together, these observations raise the possibility that distal colonic aganglionosis contributes to the perinatal death of *Rarb^R387C/R387C^* mice. Although we did not observe any change in the development of the ENS in *Rarb^R387C/+^* embryos (Fig. S2B), it is noteworthy that colonic hypoganglionosis has been reported in an RARB-RD individual heterozygous for the p.R387C variant^36^.

Mice heterozygous for either *Rarb* variant were viable but displayed significant decrease in body weight as compared to littermate controls (Fig. 1A,B). *Rarb^R387C/+^* females displayed lower bodyweight than *Rarb^L402P/+^*females (Fig. 1B), whereas no significant difference was observed between mutant males. Like RARB-RD individuals, *Rarb^R387C/+^* and *Rarb^L402P/+^*animals exhibited microphthalmia as shown by a 16 ± 2% (p < 0.001) and 17 ± 3% (p < 0.001) decrease in eyeball diameter, respectively, when compared to wild-type littermates (Fig 1C,D). In addition, pupil diameter was significantly smaller in *Rarb^R387C/+^* (decrease of 85 ± 10 %; p < 0.001) and in *Rarb*^L402P/+^ (decrease of 86 ± 10 %; p < 0.001; Fig 1C,E) mice than in controls. No sex related difference was observed. Histological analysis revealed that the eyes of E18.5 *Rarb^R387C/R387C^* embryos exhibited a hypercellular cornea, a persistent and hyperplastic primary vitreous body (PHPV) and retinal dysplasia compared to *Rarb^+/+^* embryos (Fig. S2D-I). *Rarb^R387C/+^*embryos display a milder form of PHPV without any other eye anomalies (Fig. S2D-I). Similar to *Rarb^R387C/+^* animals, the presence of PHPV has previously been reported as the only morphological eye anomaly in *Rarb^-/-^*mice^26^.

**Figure 1.**
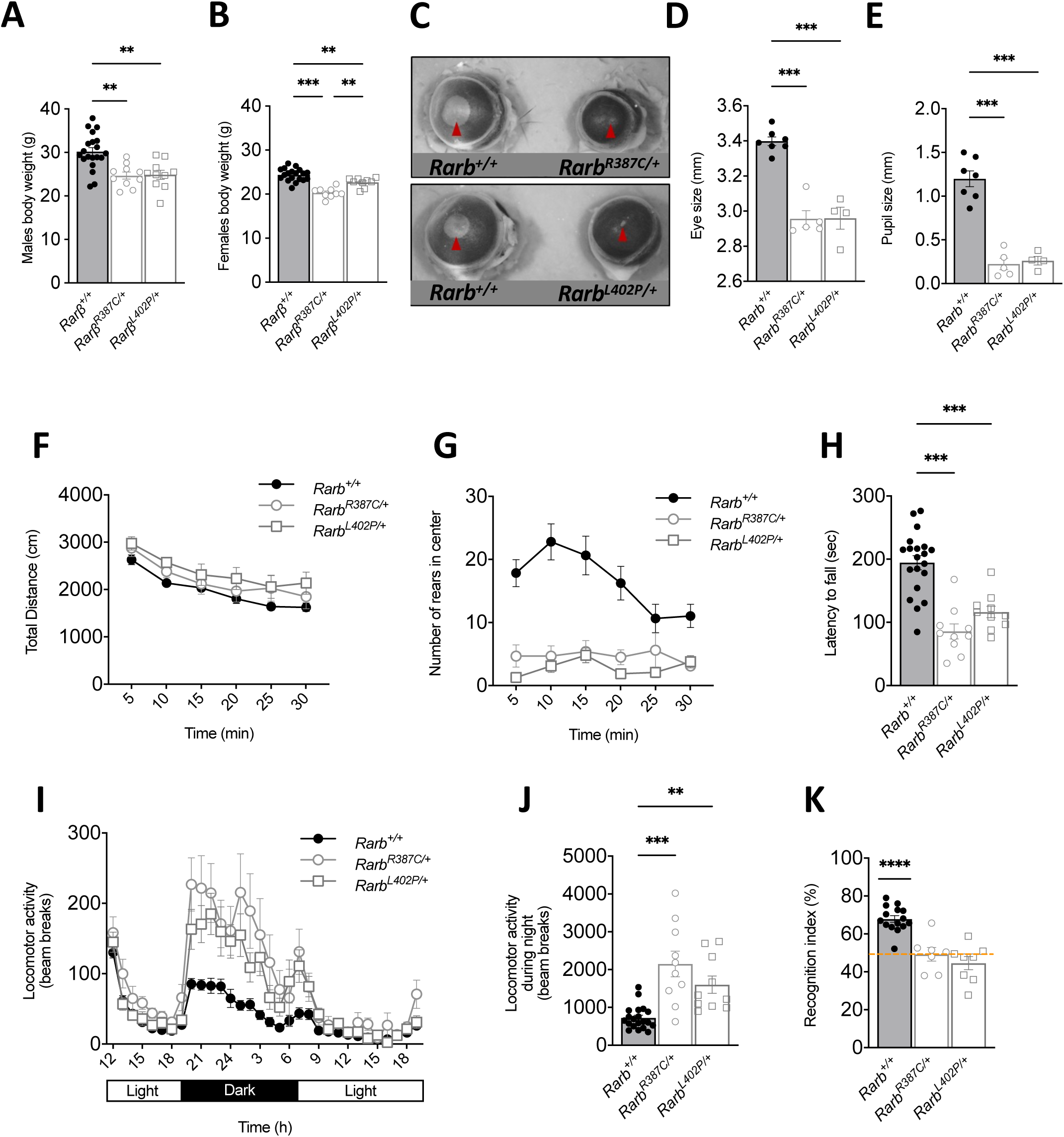
Phenotype of adult RARB-RD mice. (A,B) Bodyweight of 20-week-old males (A) and females (B). (C) Examples of eyes from 20-week-old mice. Red arrows indicate pupil. (D,E) Evaluation of eye ball size (D) and (E) pupil size. (F) Novelty induced locomotor activity in the open field (n=10 for each line). (G) Quantification of the number of rears in the center of the open field overtime. (H) Mean latency to fall from the rotating cylinder in the accelerated rotarod paradigm. (I) Spontaneous locomotor activity was measured in actimetric cages over 32 h (n=10 for all genotypes). (J) Spontaneous locomotor activity was measured in actimetric cages during 12 hours of the dark period. (K) Short-term declarative memory was assessed in novel object recognition assay. Recognition index was calculated and plotted. Orange line indicates at chance performance. Statistical differences were calculated using Bonferroni multiple comparisons as post-hoc follow-up of ANOVA and indicate: *, p < 0.05; **, p < 0.01; ***, p < 0.001. Error bars represent SEM.

Next, we investigated the motor behavior of both mouse models at 18-20 weeks of age. Although the presence of either variant did not affect the overall distance covered during 30 minutes in the open-field (Fig. 1F), both *Rarb^R387C/+^* and *Rarb^L402P/+^* mice displayed significant reduction in rearing activity which was decreased by 52 ± 13% (p < 0.001) and 27 ± 27% (p ≤ 0.05), respectively, when compared to wild-type littermates (Fig. S3A). We speculated that such reduction may reflect coordination deficits in which case rearing would be more affected in the center of the open field where mice cannot lean against walls. Indeed, the number of rears in the center of the open-field was reduced by more than 70% for both lines (Fig. 1G; S3B). This deficit did not result from reduced time spent in the center of the open field which was not significantly different between mutant mice and littermate controls (Fig. S3C). Compromised motor coordination was further supported by deficits in the accelerated rotarod test. Accordingly, latency to fall of *Rarb^R387C/+^* and *Rarb^L402P/+^* mice was decreased by 56 ± 13 % (p < 0.001) and 40 ± 10 % (p < 0.001), respectively, when compared to control *Rarb^+/+^* mice (Fig 1H). In addition, mutant mice showed an increase in the number of slips on a notched bar with *Rarb^R387C/+^* mice being more affected than *Rarb^L402P/+^* mice (Fig. S3D). It is unlikely that motor deficits observed in these tasks reflect low muscular performance as the performance of mutant mice in the grip and hanging tests was not different from that of control mice (Fig. S3E,F). Mouse models of genetic forms of dystonia typically display motor coordination deficits, including increased slips on the horizontal beam test and poor performance on the rotarod test, without any dystonic features (reviewed in^37^). The motor deficits observed in the RARB-RD mice are thus consistent with those observed in other models of dystonia.

In addition to evaluating novelty-induced activity in the open field, we also investigated spontaneous activity over a complete light/dark cycle. Globally, both mutant lines displayed normal circadian activity pattern, and their reactivity to novelty was not altered as indicated by their locomotion during the first hour of the habituation phase which was comparable to wild-type littermates (Fig. 1I). However, we observed a strong, almost 3-fold increase in locomotion during the active dark phase (between 7pm and 7am) for both *Rarb^R387C/+^*(296 ± 115%; p < 0.001) and *Rarb^L402P/+^* (221 ± 83%; p < 0.01) mice as compared to littermate controls (Fig. 1I,J).

To determine whether the mutations affect cognition, we investigated episodic memory in both mouse models using the novel object recognition test. We found that 18- to 20-week-old *Rarb^R387C/+^* and *Rarb^L402P/+^* mice displayed significant deficits in this task. As revealed by one group t-test, the performance of both mutant lines was not different from chance level of 50% at short, 3 hours, inter-trial interval (p = 0.842 for *Rarb^R387C/+^* and p = 0.166 for *Rarb^L402P/+^*), but it was significantly higher than chance level for control littermates (p < 0.001) (Fig. 1K). We excluded the possibility of a confounding effect induced by increased anxiety, as naive *Rarb^R387C/+^*, *Rarb^L402P/+^* and wild-type littermates spent a comparable amount of time in the open, anxiogenic arms of the elevated plus maze (Fig. S3G,H).

We previously reported that *Rarb^-/-^* mice display a similar profile of motor deficits as that of RARB-RD mice^24^ but without any deficits in the novel object recognition test^38^. We sought to determine whether *Rarb* haploinsufficiency also affects motor behavior. We did not observe any effect of a deletion of a single copy of *Rarb* on the number of rears in the open field, motor coordination in the rotarod or locomotion during the dark period in actimetric cages (Fig. S3I-K). In addition, performance of *Rarb^+/-^* mice was significantly higher from the chance level of 50% (p < 0.001) in the novel object recognition test, indicating that *Rarb* haploinsufficiency does not affect cognition in the context of this paradigm (Fig. S3L).

### Motor and cognitive deficits are progressive in *Rarb^R387C/+^* mice

We next sought to determine when body weight, motor and cognitive deficits first appear in *Rarb^R387C/+^* mice. We identified significant bodyweight deficits starting from 5 weeks of age in males (Fig. 2A) and 3 weeks in females (Fig. 2B). To address behavioral abnormalities, we monitored across different ages the key parameters affected in 18- to 20-week-old *Rarb^R387C/+^* mice. We found a strong tendency for *age x genotype* interaction (F (1, 20) = 3,9, p=0.06) for rearing in the central part of the arena where *Rarb^R387C/+^*mice displayed decreasing profile between 5 and 12 weeks of age in contrast to increasing number of rears in control wild-type littermates (Fig. 2C). When the performance of 18- to 20-week-old mice was included in these analyses, the age × genotype interaction was significant (F (2,30) = 3.61, p < 0.05), indicating that the onset of this deficit is at about 12 weeks of age. We next asked whether reduced rearing activity could reflect major coordination deficits which took more time to develop than finer coordination deficits which could be detected at earlier ages in other dedicated tests. Accordingly, latency to fall from the rotarod was already strongly decreased in 3 weeks-old *Rarb^R387C/+^* mice (65 ± 14% reduction as compared to *Rarb^+/+^*, p < 0.001; Fig. 2D) whereas at the same age the number of slips on the notched bar was twice as high as those scored in wild-type littermates (increase of 200 ± 102 % as compared to *Rarb^+/+^*, p < 0.01; Fig. 2E). Spontaneous locomotion during the dark period in actimetric cages also evolved differently in time for wild-type and mutant mice as revealed by significant interaction of *age x genotype* in two-way ANOVA (F (1, 11) = 7,349, p<0.05, Fig. 2F,G). Post-hoc analyses indicated that although spontaneous locomotion increased between 5 and 12 weeks of age for both *Rarb*^+/+^ and *Rarb*^R387C/+^ mice, such an increase was significantly higher for *Rarb*^R387C/+^ animals establishing 12 weeks as the onset of the hyperactivity phenotype. Finally, cognitive deficits were also age-dependent as documented by *age x genotype* interaction in two-way ANOVA (F (1, 12) = 9,404, p<0.01; Fig 5H). Accordingly, deficits observed in 18- to 20-week-old mutant mice were already present at 12 weeks (49 ± 6% for *Rarb^R387C/+^* vs. 67 ± 5% for *Rarb^+/+^*, p < 0.01), but absent at 8 weeks of age (58 ± 6% in *Rarb^R387C/+^ vs* 64 ± 7% in *Rarb^+/+^*, ns). Taken together, these observations suggest the involvement of a neurodegenerative process in *Rarb^R387C/+^* mice.

**Figure 2.**
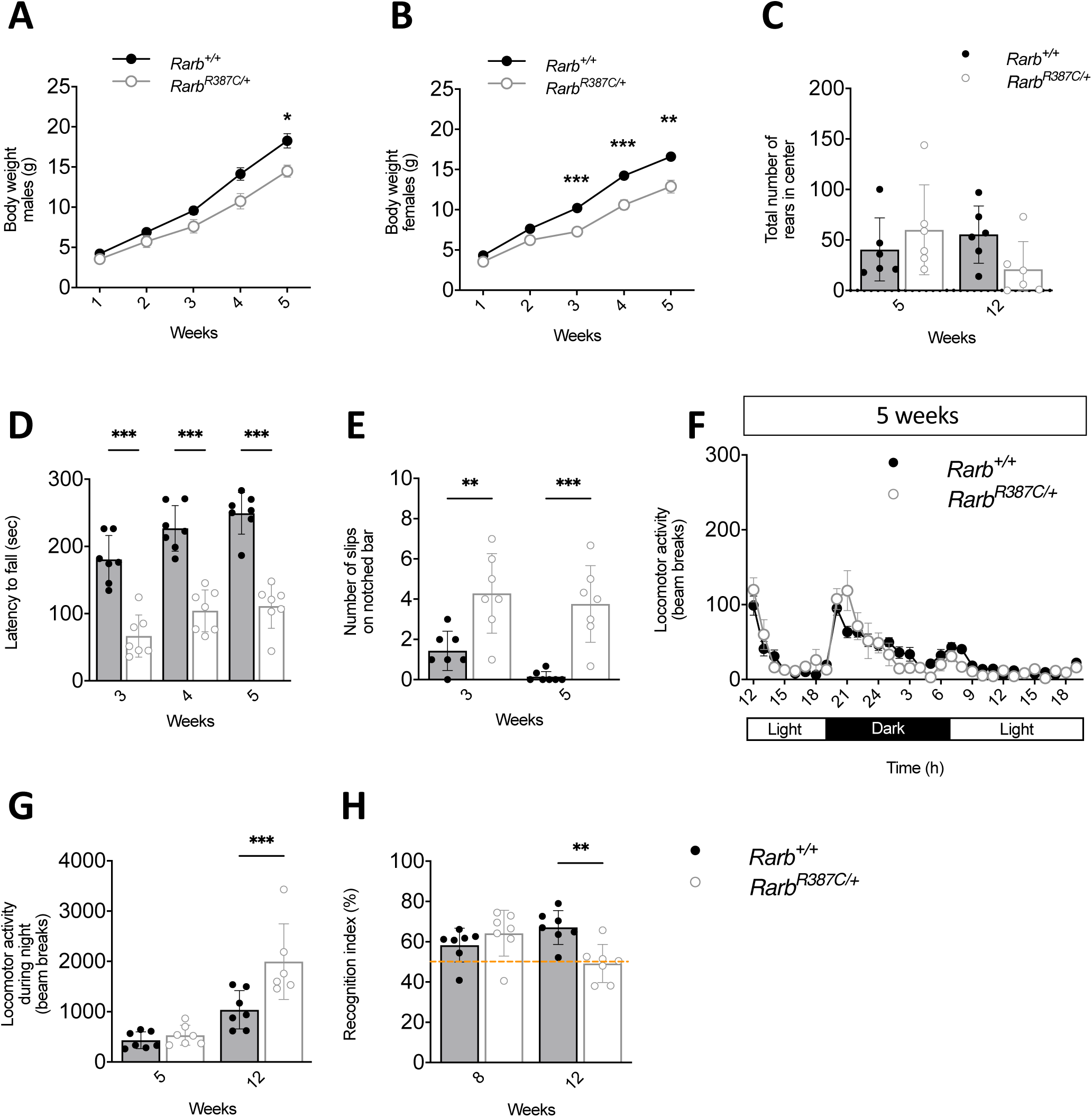
Progressive onset of symptomatology in *Rarb^R387C+/-^* mice. (A) Body weight progression between 1 and 5 weeks of age. (B) Body weight progression between 1- and 5-week-old females. (C). Quantification of the total number of rears during 30 minutes in the center of open field arena. (D) Progression of motor coordination deficits in the accelerated rotarod was expressed as mean latency to fall from the rotating cylinder. (E) Quantification of the number of slips on a 1 meter notched bar in 3- and 5-week-old mice. (F) Spontaneous locomotor activity of 5-week-old mice was measured in actimetric cages over 32 hours including 12hrs of light/dark cycle (n = 7/genotype). (G) Spontaneous locomotor activity in actimetric cages during the dark period was compared at 5 and 12 weeks of age. (H) Short-term declarative memory was assessed with novel object recognition assay at 8 and 12 weeks of age. Recognition index was calculated and plotted. Dashed orange line indicates at chance performance. Statistical differences were calculated using Bonferroni multiple comparisons as post-hoc follow-up of ANOVA analyses and indicate: *, p < 0.05; **, p < 0.01; ***, p < 0.001. Error bars represent SEM.

### RARB-RD variants affect RARB protein levels in the striatum

RARB is known for controlling its own expression by binding to regulatory elements within its promoter^39^. To study whether the RARB-RD variants affect RARB protein levels, we quantified its expression in the striatum of newborn hetero- and homozygous mice for each studied variant. RARB protein levels were significantly reduced in both heterozygous (by around 37% in *Rarb*^R387C/+^ and 53% in *Rarb*^L402P/+^) and homozygous (by around 67% in *Rarb*^R387C/R387C^ and 63% in *Rarb*^L402P/L402P^) mice when compared to wild-type littermates (Fig. 3A,B; S4A,B). The reduction of RARB expression was maintained in adult heterozygous mutants, as compared to *Rarb^+/+^* controls (Fig. 3C; S4C). To determine whether this reduction of RARB expression correlates with the behavioral deficits observed in mice carrying RARB-RD variants, we also quantified RARB expression in *Rarb^+/-^* mice which do not display any behavioral deficits (Fig. S4I-L). We observed a significant 42 ± 1% decrease of RARB. The protein was not detectable in *Rarb^-/-^* striatal samples (Fig. S4D,E). Thus, since global RARB production levels was not a determinant of behavioral deficits, we searched whether RARB-RD variants may affect nuclear localization of the protein. To this end, we quantified nuclear and extranuclear expression of p.R387C and p.L402P variants in the striatum of newborn mice homozygous for each RARB-RD variant. Using quantitative immunofluorescence, we found a 22 ± 2% (p ≤ 0.05) and 35 ± 1% (p < 0.001) reduction of nuclear RARB variants in *Rarb*^R387C/R387C^ and *Rarb*^L402P/L402P^ mice respectively, as compared to wild-type littermates (Fig. 3D,E). Cytoplasmic RARB signal was not affected by any of the variants, thus validating mostly a decrease of mutant forms of RARB without any impact on their subcellular traffic (Fig. 3D,E).

**Figure 3.**
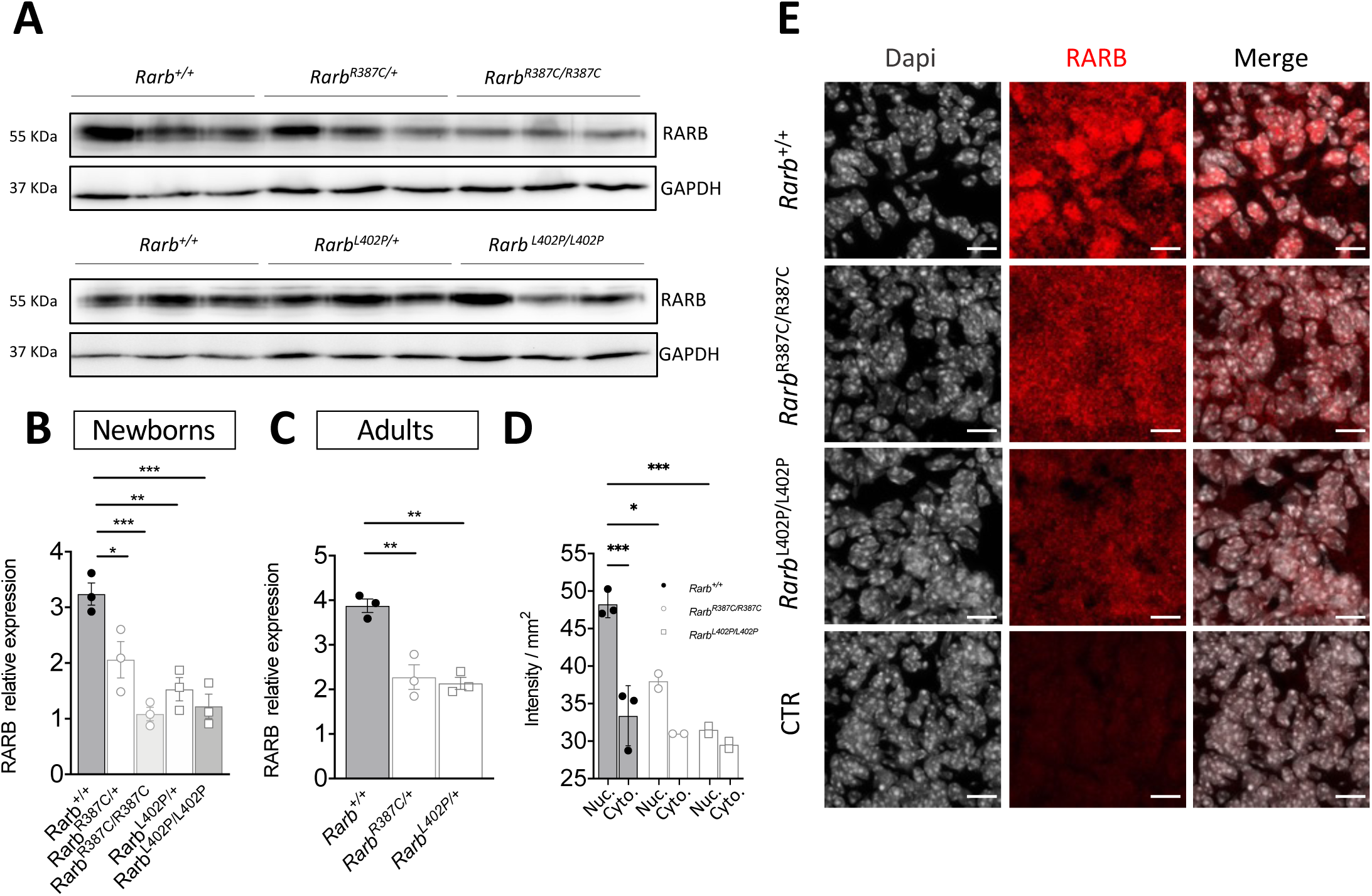
RARB production in the striatum of RARB-RD mice. (A) Western blot detection of RARB in the striatum samples from heterozygote and homozygote RARB-RD newborn mice (cropped gels, full gels are shown in Fig. S4A, S4B) and (B) quantification of RARB levels relative to those of GAPDH in these samples. (C) Western blot quantification of RARB expression levels in 20-week-old mouse striatal samples relative to those of GAPDH. Full gels are available in Fig. S4C. (D) Measurement of RARB immunofluorescence intensity per mm^2^ inside nucleus (Nuc.) or in cytoplasm (Cyto) in quantitative immunodetection of RARB as illustrated by the red signal (E) on 14µm striatal cryosections from newborn mice. CTR corresponds to control without primary antibody. Error Bar = 10µm. Statistical differences indicate: *, p < 0.05; **, p < 0.01; ***, p < 0.001. Error bars represent SEM.

### Striatal dopamine D2R signaling is compromised in RARB-RD mice

Considering that *Rarb* expression in the striatum is required for MSN development and function as well as for motor coordination^2,22^, we investigated the integrity of MSN composition and dopaminergic signaling in the dorsal striatum of adult mice carrying the RARB-RD variants. Using *in situ* hybridization to identify MSNs and their subtypes, we first observed an overall decrease of GAD67-positive GABAergic MSNs in *Rarb^R387C/+^* (15 ± 6%) and *Rarb^L402P/+^*(15 ± 3%) as compared to *Rarb*^+/+^ mice (Fig. 4A,B). Interestingly, although the number of *Drd1*-expressing cells was not affected in both mutant lines (Fig. 4A,B), the number of cells expressing *Tac1*, a specific marker of snMSNs, was decreased by 24 ± 6% in *Rarb^R387C/+^*(p < 0.001) and 19 ± 3% in *Rarb^L402P/+^* (p < 0.01) as compared to wild-type littermates (Fig. S5A,B). This observation reflects most probably a decrease of *Tac1* expression rather than the loss of *Tac1*-positive cells given the unchanged number of *Drd1*^+^ MSNs. In contrast, we observed a significant decrease in cells expressing *Drd2* (21 ± 8% *in Rarb^R387C/+^*, p ≤ 0.01 and 16 ± 5% in *Rarb^L402P/+^*, p ≤ 0.05) (Fig. 4A,B) and a comparable decrease in cells expressing *Penk*, a distinct marker of spMSNs (about 20% decrease in both models) (Fig. S5A,B). Finally, we generated adult *Rarb^R387C/+^* and *Rarb^+/+^* mice that are hemizygous for the *Drd1a-tdTomato*^40^ or *Drd2-gfp*^41,42^ reporter transgenes as highly sensitive, alternative approach to investigate the two types of MSNs. In agreement with *in situ* hybridization data, we did not observe any significant difference in the number of cells expressing the *Drd1a-tdTomato* transgene between the striatum of *Rarb^R387C/+^* and *Rarb^+/+^* mice (Fig. S6A-H) but we found a significant decrease in the number of cells expressing the *Drd2-gfp* transgene across the striatum of *Rarb^R387C/+^*mice when compared to *Rarb^+/+^* mice (Fig. S6I-P). Taken together, these results indicate that there is a decrease in the number of spMSNs, but not snMSNs, in RARB-RD mouse models.

**Figure 4.**
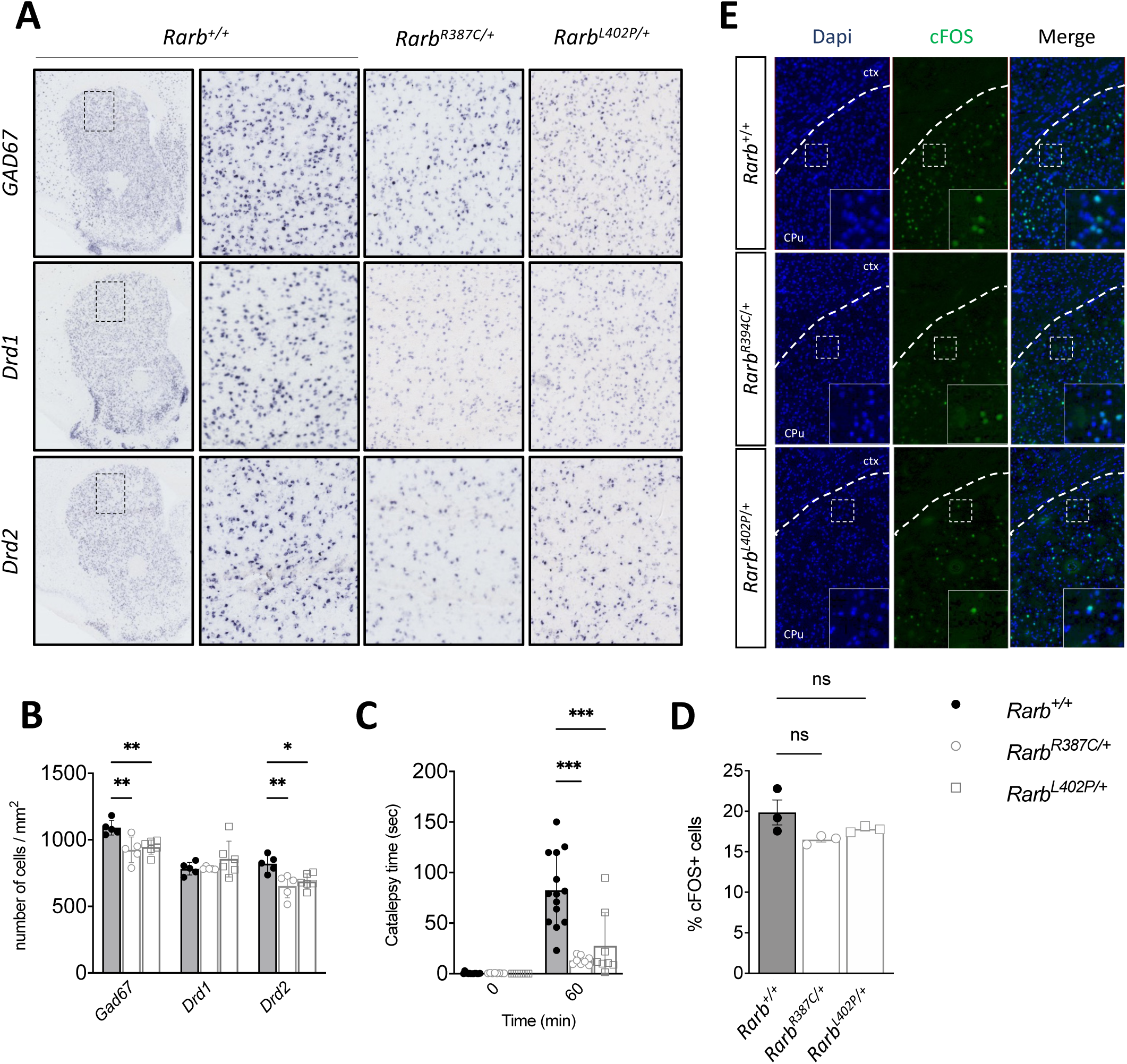
Decrease number of spMSNs and compromised spMSN signaling in RARB-RD mice. (A,B) Identification of *GAD67*-positive neurons, snMSNs (*Drd1*) and spMSNs (*Drd2*) on striatal sections from wild-type and heterozygous RARB-RD mice using *in situ* hybridization (A). Selected regions of dorsolateral striatum (see rectangles) were enlarged in adjacent panels and correspond to the areas used for counting the number of cells positive for *GAD67*, *Drd1* and *Drd2* (B). (C) Catalepsy evaluated as latency to move in the bar-test was measured before (0 minute) and 60 minutes after haloperidol (1 mg/kg) treatment. (D) Evaluation of proportion of haloperidol-induced cFOS expression (% of total number of cells) in the dorsolateral CPu 90 min after treatment. (E) Examples of immunodetection of cFOS expression in the striatum (CPu). White squares represent regions selected for illustrating at high resolution cFOS expression in CPu. Statistical differences were calculated using Bonferroni multiple comparisons as post-hoc follow-up of ANOVA analyses and indicate: *, p < 0.05; **, p < 0.01; ***, p < 0.001. Error bars represent SEM.

To further address whether the striato-pallidal pathway is functionally compromised, we challenged spMSN signaling *in vivo* by administering haloperidol, a D2R-specific antagonist, to the mutant mice and their control littermates. As expected, 1mg/kg of haloperidol efficiently induced catalepsy in wild-type mice, whereas *Rarb^R387C/+^* and *Rarb^L402P/+^* mice were much less sensitive to such treatment, displaying an 85 ± 4% (p < 0.001) and 67 ± 30% (p < 0.001) reduction in catalepsy, respectively, compared to *Rarb*^+/+^ littermates (Fig. 4C). The proportion of cFOS+ cells was comparable between all tested lines (Fig. 4D, E), indicating that some components of the D2R signaling pathway are preserved in the remaining spMSNs.

### Pramipexol-mediated modulation of D2R signalling does not rescue the behavioural phenotypes of *Rarb^R387C/+^* mice

To further investigate the hypothesis of disrupted D2R signaling in RARB-RD, we tested whether enhancing D2R signaling using pramipexol, a preferential D2R agonist, can normalize behavioral deficits in *Rarb^R387C/+^* mice. To this end, we used pramipexol at suboptimal (0.2 mg/kg) and submaximal (2 mg/kg) doses. Consistent with previous observations^43^, pramipexole induced a dose-dependent reduction of motor activity in wild-type mice when tested in the open-field, but this effect was also observed in *Rarb^R387C/+^* mice as indicated by the main effect of *treatment* (F (2,36) = 16.2, p<0.001) without any *treatment x genotype* interaction in the two-way ANOVA test (Fig. 5A). In contrast, while rearing in the central part of the arena was suppressed in *Rarb^+/+^* mice at 2mg/kg of pramipexole, no significant decrease was observed in *Rarb^R387C/+^* mice, most likely due to a floor effect in this behavior, which was already very low under both untreated and vehicle-treated conditions (compare Fig 1G and Fig. 5B). *Rarb^R387C/+^* mice were also resistant to the effects of pramipexole on locomotor coordination in the accelerated rotarod test, whereas wild-type mice displayed a dose-dependent decrease in latency to fall which is supported by significant *treatment x genotype* interaction in two-way ANOVA (F (2,36) = 6.9, p<0.01) (Fig. 5C). Similarly, whereas pramipexole dose-dependently increased spontaneous locomotion of wild-type mice in the actimetry cages it did not significantly modify the behavior of *Rarb^R387C/+^* mice (Fig. 5D). The resistance of *Rarb^R387C/+^*mice to pramipexole provides additional support for the disruption of D2R signaling in RARB-RD spMSNs but it also excludes the use of this compound as a possible symptomatic therapy in RARB-RD.

**Figure 5.**
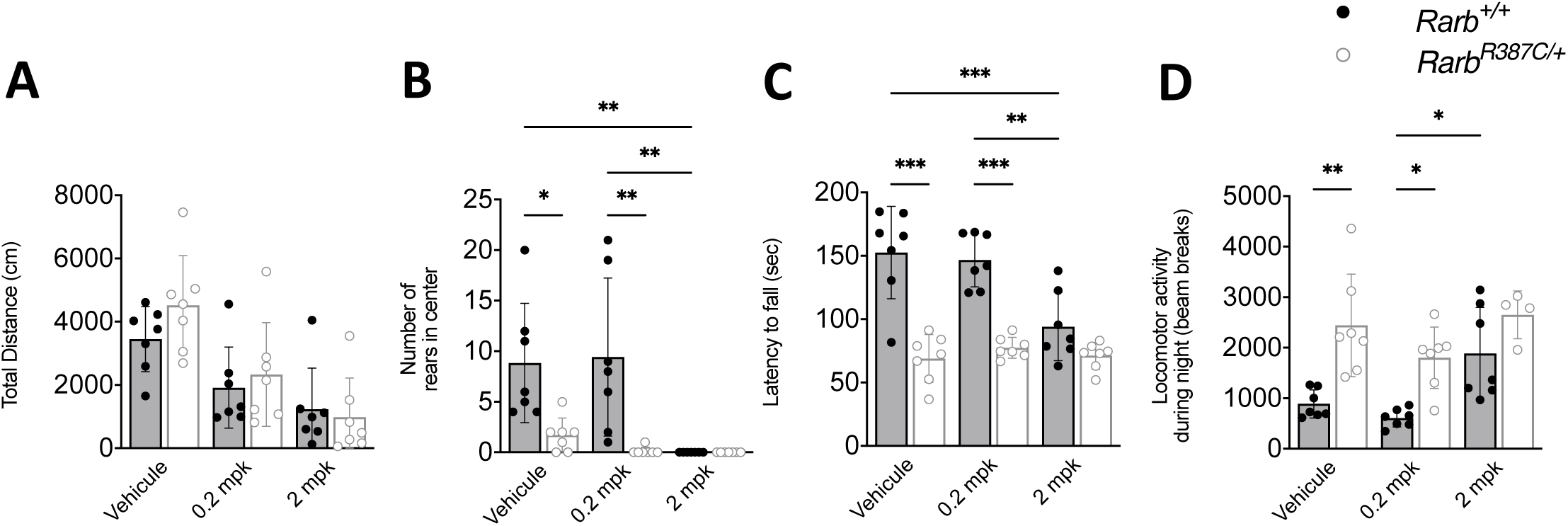
Insensitivity of RARB-RD mice to pramipexol. (A) Novelty induced locomotor activity in open field evaluated 30 minutes after intra-peritoneal injection of 0.2 mg/kg (mpk) or 2 mpk of pramipexol. (B) Quantification of the number of rears in the center of the open field 30 minutes after intra-peritoneal injection of 0.2 mpk or 2 mpk of pramipexol. (C) Motor coordination was evaluated in accelerated rotarod as mean latency to fall from the rotating cylinder 45 minutes after intra-peritoneal injection of 0.2 mpk or 2 mpk of pramipexol. (D) Long term effect of pramipexol on night spontaneous activity measured in actimetric cages. Pramipexol was injected at the onset of the dark period (7pm) after 8 hours of habituation in actimetric cages. Statistical differences were calculated using Bonferroni multiple comparisons as post-hoc follow-up of two-way ANOVA analyses and indicate: *, p < 0.05; **, p < 0.01; ***, p < 0.001. Error bars represent SEM.

### RARB-RD variants are associated with similar transcriptomic profiles in the striatum

We previously reported that p.R387C and p.L402P behave as GOF and DN variants, respectively, in cell-based transcriptional assays. We sought to determine whether these *in vitro* effects correlate with distinct transcriptional signatures *in vivo* by performing bulk RNAseq to compare the dorsal striatal transcriptomes of two-month-old *Rarb^R387C/+^*, *Rarb^L402P/+^*, *Rarb^-/-^*and *Rarb^+/-^*males with those of paired control littermates. We found 2364 differentially expressed genes (DEGs) in the striatum of *Rarb^R387C/+^*mice and 2520 in *Rarb^L402P/+^*mice (Fig. 6A,B; Table S1). In contrast, we observed 800 DEGs in the striatum of *Rarb^-/-^* mice, a much smaller number of genes than in RARB-RD mice, whereas the number of DEGs in *Rarb^+/-^* mice was only 99 (Fig. 6A,B; Table S1). These results suggest that heterozygous RARB-RD variants have a more profound impact on the striatal transcriptome than the null allele whether it is found in the homozygous or heterozygous state.

**Figure 6.**
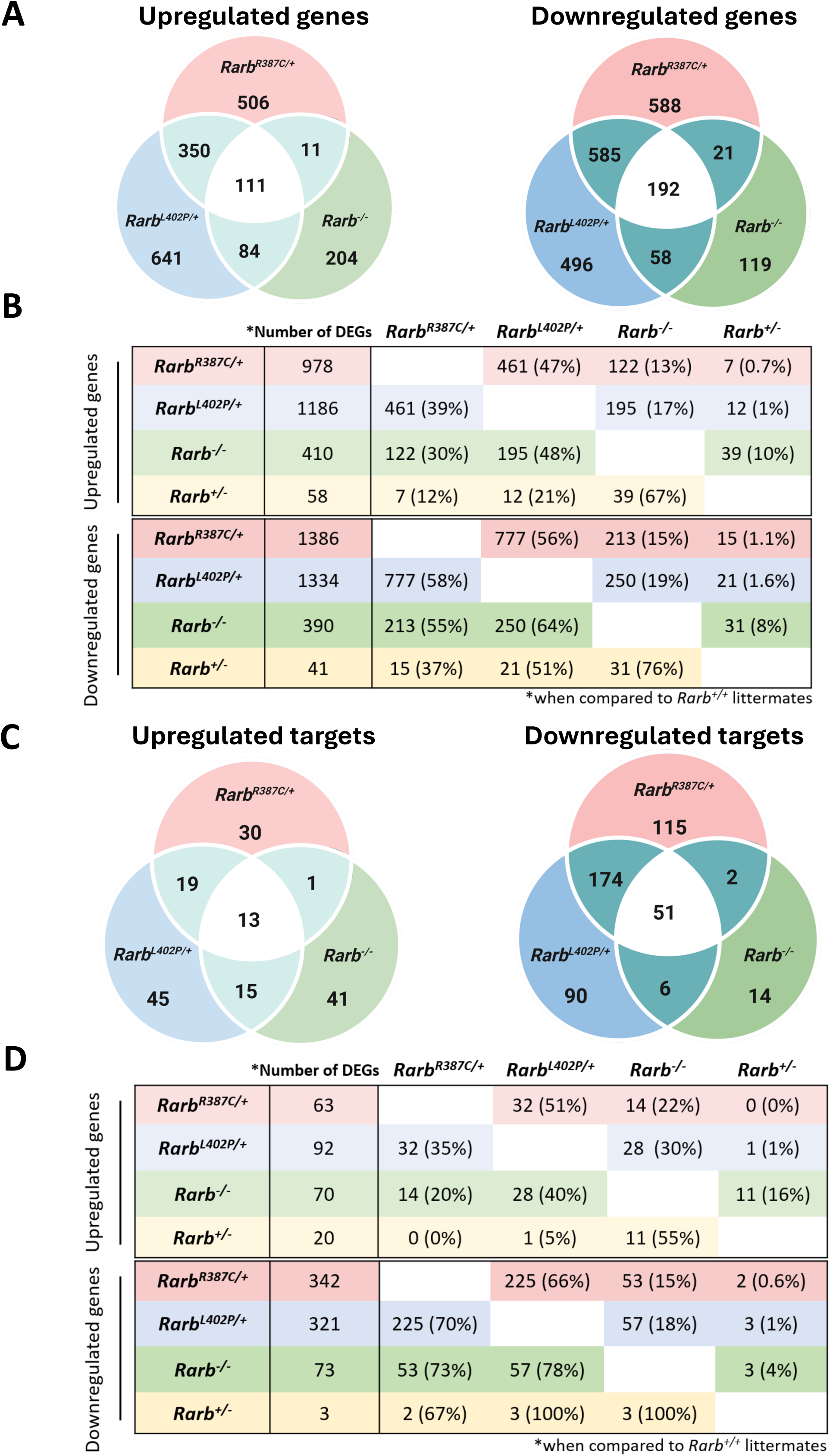
Striatal transcriptomes of *Rarb* mutant mice. (A) Overlap of upregulated (left panel) and downregulated (right panel) genes in *Rarb*^R387C/+^, *Rarb*^L402P/+^, and *Rarb*^−/−^ mice compared to their respective *Rarb*^+/+^ littermates. DEGs were defined by a FDR-corrected Q-value cutoff of 0.05. (B) Number of DEGs in the striatum of mice with a given genotype that are also upregulated or downregulated in mice with the other genotypes. (C) Overlap of upregulated (left panel) and downregulated (right panel) genes encoding putative direct targets of RARB in *Rarb*^R387C/+^, *Rarb*^L402P/+^, and *Rarb*^−/−^ mice compared to their respective *Rarb*^+/+^ littermates. DEGs were defined by a FDR-corrected Q-value cutoff of 0.05. (D) Number of DEGs encoding direct targets of RARB in the striatum of mice with a given genotype that are also upregulated or downregulated in mice with the other genotypes. The number of mice used for these experiments is as follow: *Rarb^R387C/+^*(n=5) and *Rarb^+/+^* (n=5); *Rarb^L402P/+^* (n=4) and *Rarb^+/+^* (n=5); *Rarb^-/-^* (n=4), *Rarb^+/-^* (n=4) and *Rarb^+/+^* (n=4). Only males were used for these experiments.

We next compared DEGs across *Rarb^R387C/+^*, *Rarb^L402P/+^*and *Rarb^-/-^* mice. We found a significant overlap in down- and up-regulated genes among these lines (Fig. 6A,B; S7). Moreover, DEGs were significantly enriched for a similar subset of KEGG and/or Reactome functional terms, including a number of neuronal signaling and synaptic transmission pathways (Fig. 7A; S8; Table S1). We also found that DEGs were significantly enriched in all the mutant lines for putative direct targets of RARB that were identified in ChIP-seq experiments performed on striatal samples from adult mice (Table S1)^39^. The majority of the DEGs encoding these targets were downregulated in mutant mice (*Rarb^R387C/+^*: 342/405, 84%; *Rarb^L402P/+^*: 321/413, 78%; *Rarb^-/-^*: 73/143, 51%) and approximately 2/3 of them were downregulated in both *Rarb^R387C/+^*and *Rarb^L402P/+^* mice (Fig. 6C,D). Downregulated genes encoding putative targets in *Rarb^R387C/+^* or in *Rarb^L402P/+^*mice were significantly enriched for a number of the same KEGG and Reactome functional terms as all DEGs, suggesting that these targets play a major role in the disruption of the striatal transcriptome by RARB-RD variants (Fig. 7A; S8). The observation that the majority of DEGs encoding RARB targets are downregulated in *Rarb^R387C/+^*, *Rarb^L402P/+^* and *Rarb^-/-^* mice, and that these sets extensively overlap suggests that both the p.R387C and p.L402P variants reduce RARB function and disrupt its signaling through a shared mechanism.

**Figure 7.**
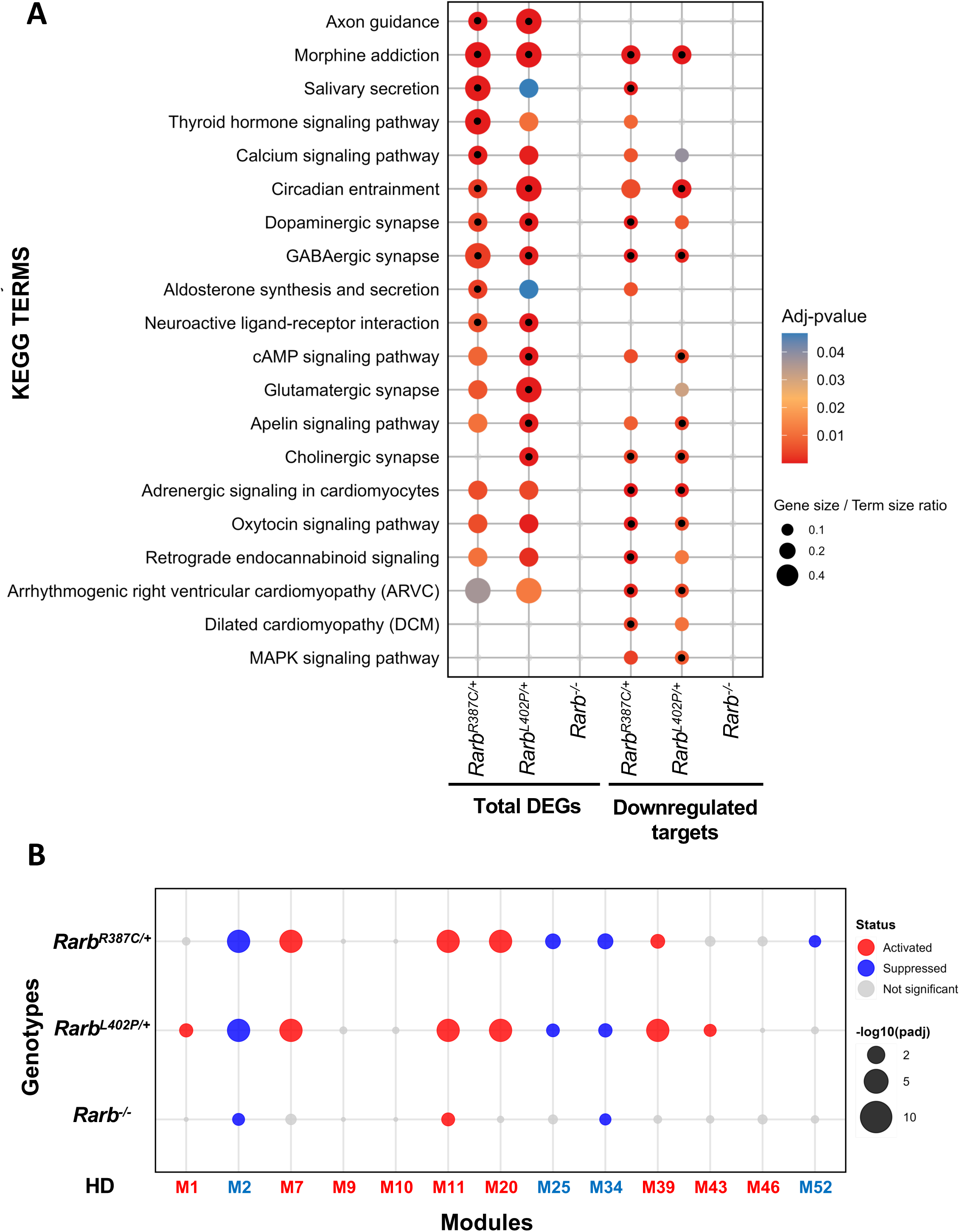
Pathways and gene sets enrichment in the transcriptomes of RARB-RD and *Rarb^-/-^* mice. (A) KEGG analysis of all DEGs or of downregulated genes encoding RARB targets in *Rarb^R387C/+^*, *Rarb^L402P/+^* or *Rarb^-/-^* mice, indicating significant enrichment of similar functional terms in RARB-RD mice. Adjusted p-values (p < 0.05) are indicated by the color code and gene/term ratio by the size of the dots. The top 10 significant pathways for each line are indicated by the black dots within the colored ones. (B) Dot-plot illustrating gene set enrichment of DEGs from *Rarb* mutant mice in the 13 gene expression modules correlated with HD CAG repeat length. Adjusted p-values (p < 0.05) are indicated by the size of the dots. For *Rarb* mutants, downregulated modules are represented by blue dots and upregulated modules by red dots whereas for HD mice, downregulated modules are shown in blue and upregulated modules in red at the bottom of the plot. The number of mice used for these experiments is as follow: *Rarb^R387C/+^* (n=5) and *Rarb^+/+^* (n=5); *Rarb^L402P/+^* (n=4) and *Rarb^+/+^* (n=5); *Rarb^-/-^* (n=4), *Rarb^+/-^* (n=4) and *Rarb^+/+^* (n=4). Only males were used for these experiments.

By performing *in vivo* RNAi-based and single-guide RNA CRISPR-based screens, Wertz et al (2020) identified 72 postnatal neuronal essential genes whose expression is enriched in the striatum when compared to the rest of the brain^44^. It is noteworthy that these genes are significantly enriched for direct targets of RARB^44^. GSEA revealed that DEGs in *Rarb^R387C/+^* and *Rarb^L402P/+^* mice, but not in *Rarb^-/-^* mice, were significantly enriched for these genes (Table S1). Among the essential genes that were differentially regulated in RARB-RD mice, the vast majority were downregulated (38/42 (90%) in *Rarb^R387C/+^* and 39/42 (93%) in *Rarb^L402P/+^*mice) (Table S1). Overall, these observations raise the possibility that RARB-RD variants affect spMSN survival by disrupting the expression of these neuronal essential genes.

The expression of the striatal-enriched essential neuronal genes identified by Wertz et al. (2020) is also reduced in the striatum of mouse models of HD^44^. Like RARB-RD, HD is characterized by progressive motor deficits and a decrease of the number of spMSNs without any change in the number of snMSNs^45^. To further investigate the overlap between the transcriptomes associated with RARB-RD and HD, we examined whether DEGs in the *Rarb* mutant lines were enriched in striatal gene modules that have been associated with CAG repeat length and disease progression in a comprehensive longitudinal bulk RNA-seq study of HD allelic series of mouse models (Table S1)^46^. Remarkably, we found that DEGs in RARB-RD mice were significantly enriched in modules (M2, M25, M34) that had a strong negative correlation with CAG repeat length (i.e. an overall downregulation of genes in these modules correlated with earlier onset of disease associated with increasing size of CAG repeats). Like in HD mouse models, genes contributing to these modules were also mostly downregulated in RARB-RD mice (Fig. 7B). M2 is enriched in cAMP-signaling, post-synaptic density proteins and striatum identity genes, M25 in glutamate receptor signaling and M34 in transcriptional factors^46^. DEGs in RARB-RD mice were also significantly enriched in four modules that are negatively associated with CAG repeat length, and represent mostly upregulated genes both in RARB-RD and HD mice (M7, M11, M20, M39) (Fig. 7B). M20 is enriched for p53 signaling, cell division and protocadherin genes, M7 and M39 are involved in stress responses, including cell death (M7) and DNA damage repair (M39) and M11 appears to be glial-related^46^. DEGs in *Rarb^-/-^* mice were enriched in only three of these modules (M2, M34 and M11) (Fig. 7B). Downregulated modules included a larger proportion of putative targets of RARB (between 25 and 37% for M2 and M25 in RARB-RD mice) than upregulated modules (between 1 and 10%) (Table S1). One possible interpretation for these results is that the reduced expression of RARB targets initiates RARB-RD by downregulating genes in M2, M25, and M34 whereas the upregulation of genes in M7, M11, M20, and M39 is involved in a later step in the pathological process.

## Discussion

We previously reported that dominant mutations in *RARB* cause a complex disorder, RARB-RD, which is characterized by birth defects, global developmental delay and dystonia. In order to gain insight into the pathogenesis of this disorder, we generated mice carrying p.R387C or p.L402P; the former variant is found in more than 30% of RARB-RD individuals whereas the latter one was identified in multiple unrelated RARB-RD individuals. We found that these models recapitulated the RARB-RD phenotype as they both displayed microphthalmia as well as RARB-RD-like behavioral impairment. For instance, both lines were similarly affected by reduced motor coordination, hyperactivity during the active period of the light/dark cycle and cognitive deficits in the novel object recognition test. Importantly, all these behavioral abnormalities with the exception of one of the measures of motor coordination, the latency to fall in the rotarod, were age-dependent, setting off between 8 and 12 weeks of age. *Rarb^L402P/+^* mice displayed a similar but a weaker phenotype than *Rarb^R387C/+^* mice as indicated by milder body weight reduction and coordination deficits in the notched bar test. These data suggest potential genotype-phenotype correlations that may have prognostic value for patient care and future design of therapeutic approaches.

Not only do the p.R387C and p.L402P variants cause similar phenotypes, but they also both reduce the number and signaling activity of spMSNs - without affecting the number of snMSNs -, a hallmark of postnatal *Rarb* loss-of-function in the striatum^22^. Compromised D2R signaling in spMSNs may lie at the core of RARB-RD pathophysiology, as it could lead to an imbalance between the direct and indirect striatal output pathways, an imbalance thought to underlie several basal ganglia disorders^47^. However, how *Rarb* variants affect D2R signaling activity is not clear. An absence of about 20% of spMSNs could contribute to such a deficit, but considering the remarkable resistance of *Rarb^R387C/+^*and *Rarb^L402P/+^* mice to haloperidol- and pramipexol-induced effects, we speculate that other mechanisms are involved. *Drd2* expression is directly regulated by RARB^24,39^, so we could postulate that its decreased expression or hypoactivity in the remaining *Rarb^R387C/+^* and *Rarb^L402P/+^* spMSNs, can further compromise signaling of the indirect pathway. However, this possibility appears unlikely as haloperidol induced cFOS expression in a comparable proportion of cells in the striatum of *Rarb^+/+^*, *Rarb^R387C/+^* and *Rarb^L402P/+^*mice, indicating that a sufficient number of D2 receptors remains in spMSNs of mutant mice and that their intracellular signaling is functional enough to induce cFOS expression. Taken together, our results suggest that RARB-RD variants do not specifically affect D2R activity in the remaining spMSNs, but that they may disrupt downstream processes that are distinct from those involved in the induction of cFOS expression.

Although both RARB-RD variants result in similar developmental anomalies—such as microphthalmia and delayed weight gain—as well as behavioral and dopaminergic deficits resembling those observed in *Rarb^-/-^* mice^22^, several observations suggest that RARB-RD variants do not cause *RARB* haploinsufficiency. First, RARB-RD homozygous mice display a more severe phenotype than mice homozygous for a null allele, as illustrated by the neonatal lethality, colonic aganglionosis and severe eye defects observed in the former but not in the latter. Second, RARB-RD heterozygous mice are phenotypically more affected than mice heterozygous or homozygous for a null allele. For instance, *Rarb^R387C/+^*and *Rarb^L402P/+^* mice, but not *Rarb^-/-^* mice, show cognitive deficits in the novel object recognition test whereas *Rarb^+/-^*mice do not display any motor or cognitive deficits. This conclusion is further substantiated by the observation that the striatal transcriptome is much more perturbed in *Rarb^R387C/+^* and *Rarb^L402P/+^* mice than in *Rarb^-/-^* mice whereas inactivation of one *Rarb* allele has only a minimal impact on gene expression in the striatum. Taken together, these observations strongly suggest that RARB-RD variants do not cause haploinsufficiency, but rather lead to the disorder by conferring a specific property to the RARB protein. Two families with bi-allelic truncating variants in *RARB* have been reported^21,48^. These families exhibited features consistent with RARB-RD; however, their neurodevelopmental outcomes were not described. Although our results suggest that bi-allelic loss-of-function variants may lead to a less severe phenotype than dominant RARB-RD variants, further studies are needed to delineate the clinical impact of this genotype.

Previous studies have reported that RARB-RD variants can be classified based on their ability to induce GOF or DN effects in response to the ligand (1µM) in *in vitro* transcriptional assays^19^. For instance, p.R387C has been shown to behave as a GOF variant, increasing the response to retinoic acid by more than 2-fold, whereas p.L402P behaves as a DN variant in the context of these assays. In contrast, our *in vivo* transcriptional data indicate that these variants and the null allele affect the expression of a large subset of the same genes. Moreover, we observed a significant enrichment of putative direct targets of RARB - primarily downregulated - among the DEGs in both RARB-RD models and *Rarb^-/-^* mice, leading to disruption of the same subset of pathways across these models. Taken together, these results strongly suggest that p.R387C, p.L402P and possibly most RARB-RD variants behave as DN variants *in vivo*. The reason for the discrepancy between the *in vitro* and *in vivo* effects of p.R387C remains unclear. Retinoids appear to regulate gene expression *in vivo* at low nanomolar concentrations^49–51^ whereas the transcriptional responses of RARB-RD variants were studied *in vitro* at micromolar concentrations of retinoids^19^. It is possible that p.R387C behaves differently depending on retinoid concentrations.

Our study has important implications for the development of strategies to treat RARB-RD individuals. The progressive nature of the motor abnormalities in *Rarb^R387C/+^* and *Rarb^L402P/+^* mice is reminiscent of the worsening of dystonia observed in some RARB-RD individuals. Moreover, we have previously shown that abolishing *Rarb* function in the striatum of adult mice is sufficient to induce coordination deficits, loss of *Drd2*-expressing cells and decrease of D2R signaling^22,39^. These observations point to a window of opportunity for therapeutic interventions early in the course of the disorder, involving the restoration of RARB signaling to prevent further progression of RARB-RD. Finally, we found that heterozygosity for *Rarb* null mutation appears to be virtually silent in mice. Similarly, preliminary observations suggest that individuals heterozygous for null alleles of *RARB* can exhibit developmental eye defects but they do not appear to display any developmental or neurological impairment^19^. Therefore, knocking-down RARB-RD allele or its product could represent a viable strategy to treat affected individuals. The RARB-RD models that we have generated represent invaluable tools for the development and validation of such therapeutic strategies.

We have found that RARB-RD variants affect the striatal expression of the same subsets of genes as in HD, including striatal-enriched neuronal essential genes and gene modules correlating with HD CAG repeat length. Other observations support mechanistic commonalities between RARB-RD and HD. For instance, we have previously shown that the expression of the *Rarb* transcript and protein is decreased in the striatum of a mouse model of HD and that the RARB protein is associated with striatal HD-related aggregates in these mice^39^. Furthermore, a significant subset of genes differentially expressed in the striatum of HD patients are putative direct targets of RARB^39^. Finally, decreasing *Rarb* gene dosage by 50 % accelerates the HD-like motor deficits and the transcriptional changes associated with the progression of these deficits in the R6/1 HD mouse model^28^. Taken together, these observations suggest that the pathogenesis of RARB-RD and HD involves a common neurodegenerative component. One possibility is that the inclusion of RARB in HD-related aggregates impairs its function, thereby contributing to the progression of HD. These results raise the possibility that successful therapeutic strategies for one of these disorders could be repurposed to treat the other.

## Acknowledgements

The *Rarb*^em1(L402P)Ics^ mutant mouse line was established at the Institut Clinique de la Souris – PHENOMIN-ICS (http://www.phenomin.fr) within the framework of the PHENOMIN 2020 call. We are grateful to James Waldron for genotyping mice for the Michaud laboratory.

## Supplementary materials

Supplementary material is available at Brain online

## Data Availability

The datasets generated and/or analyzed in this study will be provided from the corresponding author upon reasonable request.

## Supplementary Material

### Supplementary Material and Methods

#### Open-field test

Mice were tested in automated open field arenas (44 x 44 x 18 cm) made of PVC with a black floor and translucent walls and equipped with infrared sensors (Panlab, France). Briefly, mice were individually placed in the periphery of the open field and allowed to explore the apparatus freely for 30 min. The distance travelled, the number of rears and time spent in the central and peripheral areas were automatically recorded over the test session. The test was performed in a room homogeneously illuminated at 150 Lux.

#### Novel object recognition

The object-recognition task is based on the natural tendency of rodents to explore a novel object/environment and to compare it with a familiar one. The test was performed in an open-field arenas described above (Panlab, France). The objects to be discriminated were a glass marble (2.5-cm diameter) and a die (2 cm). Each animal was first habituated to the open-field arena for 45 min. The next day, mice were submitted to a 10-min acquisition trial (first trial), during which they were placed in the corresponding open-field arena in the presence of an object A (marble or die) placed at the upper left corner, 10 cm from the side walls. The time taken by the animal to explore the object A (when the animal’s snout was directed toward the object at a distance of 2 cm or less) was manually recorded. Mice that explored object A for <4 sec were excluded. Animals were then tested in the 10-min retention test (second trial) that occurred 3 hours after the acquisition trial. During the retention test, the object A was remained in the same position (the upper left corner) while another object B (marble or die, different from object A) was placed at the upper right corner of the openfield and the times tA, and tB taken by the animal to explore the two respective objects were manually recorded. The recognition index (RI) was defined as [tB/ (tA + tB)] x 100. The type of the objects and their location during the acquisition and retention phase were counterbalanced across animals. To prevent olfactory cues, objects and arena were wiped with ethanol after each test, and forceps were used to place the objects in the arena.

#### Rotarod

An accelerated (4–40 rpm in 5 min) rotarod (Bioseb, France) was used for all experiments. Each test consisted of three trials separated by 15 min recovery intervals. The latency time to fall from the rotarod was recorded. Before the first trial mice were allowed to stay on the rotarod for about 30s to habituate before starting the acceleration phase.

#### Spontaneous locomotion in actimetric cages

Spontaneous locomotor activity was measured in actimetric cages (Immetronic, Pessac, France) during 32h (11 am until 7 pm the next day). Briefly, each mouse was placed in an individual cage equipped with infra-red photo beam cells to measure horizontal movements. During the experiment mice had free access to food and water.

#### Grip test

The Grip test measures the maximal muscular strength (g) using a grid-connected isometric dynamometer (Bioseb). Mice were allowed to grip the grid with their four paws and were then pulled backward until they released it. Each mouse was submitted to 3 consecutive trials and the maximal strength developed by the mouse before releasing the grid was measured. 3-trials-averaged value was normalized to the body weight of each animal.

#### Hanging test

The hanging test measures the muscular strength using an inverted grid and placed at 50 cm above the ground. Mice are hanging on this grid and the latency to fall is measured. The test is composed of 3 trials of maximum 1 min. Averaged values of 3 trials is considered as hanging time.

#### Notched bar

Motor coordination was tested under 100-lux lighting on a 2 cm-wide and 100 cm-long natural wooden piece notched bar comprising 26 platforms of 2 cm spaced by 27 gaps of 2 cm. Animals had to cross the whole notched bar twice for training and 3 times for the test. Every instance of a back paw going through the gap was considered an error (slip). The 3-trials-averaged number of slips was quantified.

#### Elevated plus maze

The apparatus composed of 2 open arms and 2 closed arms was positioned at 50 cm above the ground. Each mouse was placed in the center of the maze with its head facing one of the closed arms. Anxiety was measured by quantifying the latency to enter the open arm and the percentage of time spent in the open arms during 5 minutes of the testing.

#### Haloperidol-induced catalepsy

The bar test was used to examine haloperidol-induced catalepsy. During the test, both forelimbs of a mouse were placed on a horizontal bar 4 cm above the ground. Each animal underwent one session of the test (baseline/spontaneous catalepsy) prior to haloperidol treatment and 60 min after haloperidol (IP) treatment. The time of immobility before stepping down the bar was considered as catalepsy score.

#### Antibodies used in immunohistochemistry and western blot experiments

| **List of Antibodies** |  |  |  |
| --- | --- | --- | --- |
| **Primary Antibodies** | **Company** | **Catalog number** | **Dilution** |
| SOX10 | R&D Systems | AF2864 | 1:500 |
| RET | R&D Systems | AF482 | 1:500 |
| cFOS | Antibodies-online | ABIN4285633 | 1:500 |
| RFP | Rockland | 600-401-379S | 1:1000 |
| GFP | Millipore | AB16901 | 1:2000 |
| NeuN | Millipore | MAB377 | 1:1000 |
| RARB | Santa Cruz Biotechnology | C-19, sc552 | 1:500 |
| GAPDH | USCN | CAB932Hu22 | 1:5000 |
| **Secondary Antibodies** |  |  |  |
| D@Mouse | Invitrogen | A-21121 | 1:500 |
| D@Rabbit | Invitrogen | A-31572 | 1:500 |
| G@Rabbit | Invitrogen | A21430 | 1:1000 |
| G@Chicken | Abcam | ab150169 | 1:1000 |
| G@Mouse | Invitrogen | A21053 | 1:1000 |
| G@Rabbit-HRP | Jackson ImmunoResearch | 111 035 44 | 1:5000 |
| G@Mouse-HRP | Invitrogen | G-21040 | 1:5000 |

### Supplementary Figures

**Fig. S1.**
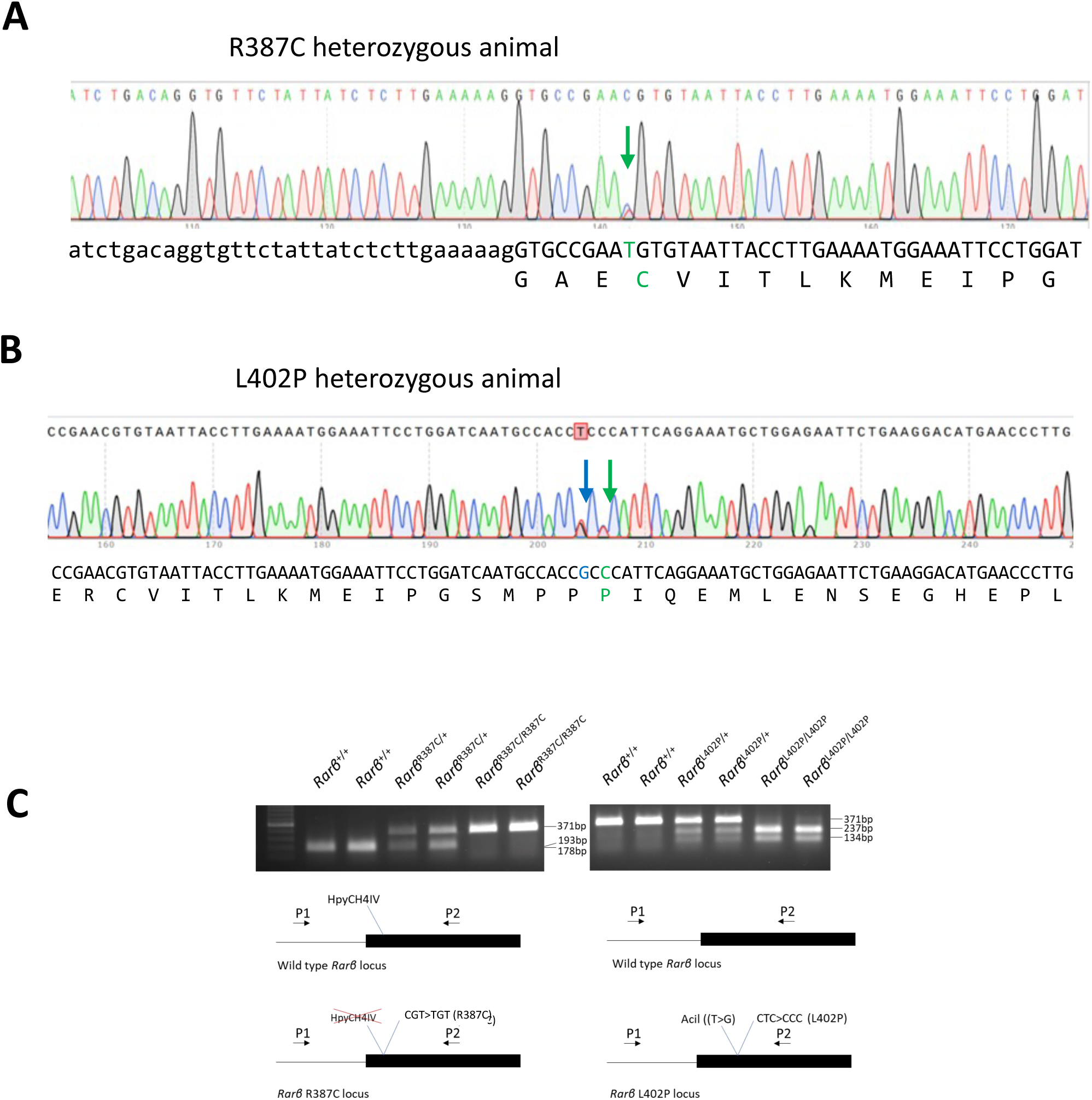
Generation of mutant mouse lines. (A, B) Chromatogram obtained by Sanger sequencing of PCR products on heterozygous (A) R387C and (B) L402P mouse. (C) Cartoon representation of WT and mutant loci and their genotyping for *the p.R387C* (left panel) and *p.L402P* (right panel) variants. In the case of p.R387C, the variant abolishes a HpyCH4IV restriction site. PCR performed with primers flanking the variant amplifies a 371bp fragment which can be cleaved by HpyCH4IV to generate 178bp and 193bp fragments from the wild-type but not the mutant allele (Fig. S1C left panel). In the case of p.L402P, a silent substitution (NM_011243.2:c.1T>G) was introduced in the preceding proline codon leading to the formation of a new Acil restriction site. PCR performed using the same primers as for p.R387C amplifies a 371 bp fragment, which can be cleaved by Acil enzyme to generate 134bp and 237bp fragments from the p.L402P but not from the wild-type allele (Fig. S1C right panel).

**Figure S2.**
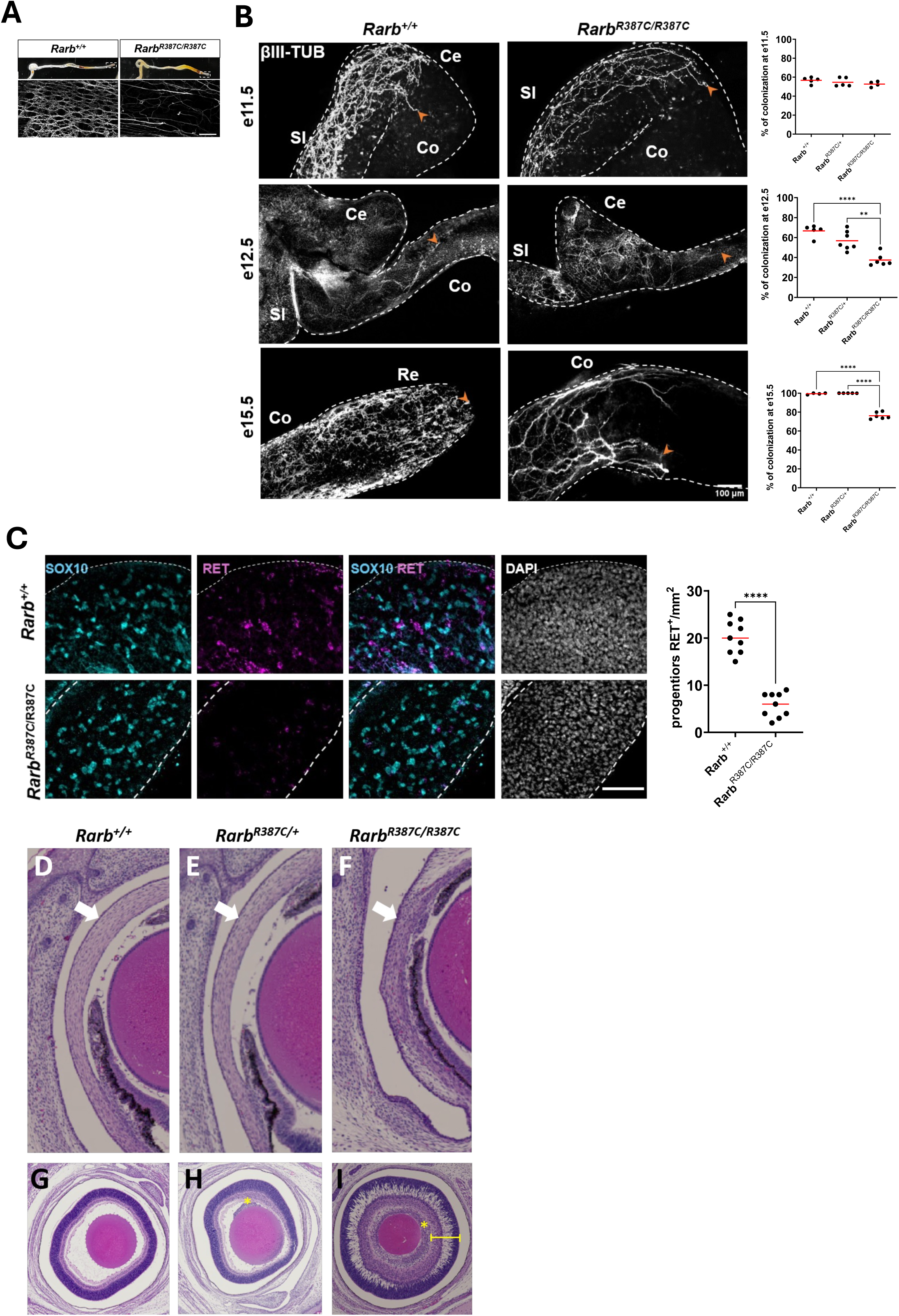
Developmental defects associated with the p.(R387C) variant. (A-C) Characterization of enteric nervous system development in *Rarb^R387C/R387C^*, *Rarb*^R387C*/+*^ and *Rarb^+/+^* embryos and mice. (A) Brightfield view of colon from *Rarb^R387C/R387C^* newborn mice shows distal blockage (arrow) and accumulation of fecal matter (asterisks), while βIII-tubulin immunofluorescence staining of distal colon (dashed boxes in upper panels) shows only a few nerve fibers in mutant mice instead of the dense networks of interconnected neural ganglia normally seen in controls. Scale bar: 500 μm. (B) βIII-tubulin immunofluorescence staining of the developing gut of E11.5, E12.5 and E15.5 *Rarb^R387C/R387C^*, Rarb^R387C*/+*^ and *Rarb^+/+^* embryos. Representative images of *Rarb^R387C/R387C^* and *Rarb^+/+^* samples are shown for each stage. Staining quantification indicates significant decrease in the extent of colonization of the colon by enteric neural progenitors at E12.5 and E15.5 in *Rarb^R387C/R387C^* when compared to *Rarb*^R387C*/+*^ and *Rarb^+/+^* embryos (One-way ANOVA with Sidak’s post hoc test, **, p < 0.01; ***, p < 0.001, ****p<0.0001; Error bars represent SEM; n = 4-7 animals per group). Scale bar: 100 μm. SI: small intestine, Co: colon, Re: rectum and Ce: cecum. (C) Double immunofluorescence staining of SOX10+ (cyan) and RET+ (pink) enteric neural progenitors in the distal foregut of E12.5 *Rarb^R387C/R387C^* and *Rarb^+/+^* embryos, showing a specific decrease of the RET signal in the mutants when compared to controls. (Student’s *t*-test, ****p<0.0001 Error bars represent SEM; n = 3 animals per group, 3 fields of view per sample). Scale bar: 100 μm. (D-I) Hematoxylin-stained sagittal sections of the eyes of E18.5 paraformaldehyde-fixed *Rarb^+/+^* (D,G), *Rarb^R387C/+^* (E, H), and *Rarb^R387C/R387C^* (F,I) embryos. The cornea (white arrow) appears normal in *Rarb^+/+^* (D) and *Rarb^R387C/+^* (E) embryos but hypercellular in *Rarb^R387C/R387C^* embryos (F). *Rarb^R387C/+^* (H) and *Rarb^R387C/R387C^* (I) embryos display persistent hyperplastic primary vitreous (yellow asterisk and arrow) whereas *Rarb^R387C/R387C^* (I) embryos also ehxibit diffuse irregularity of retinal architecture (bracket).

**Figure S3.**
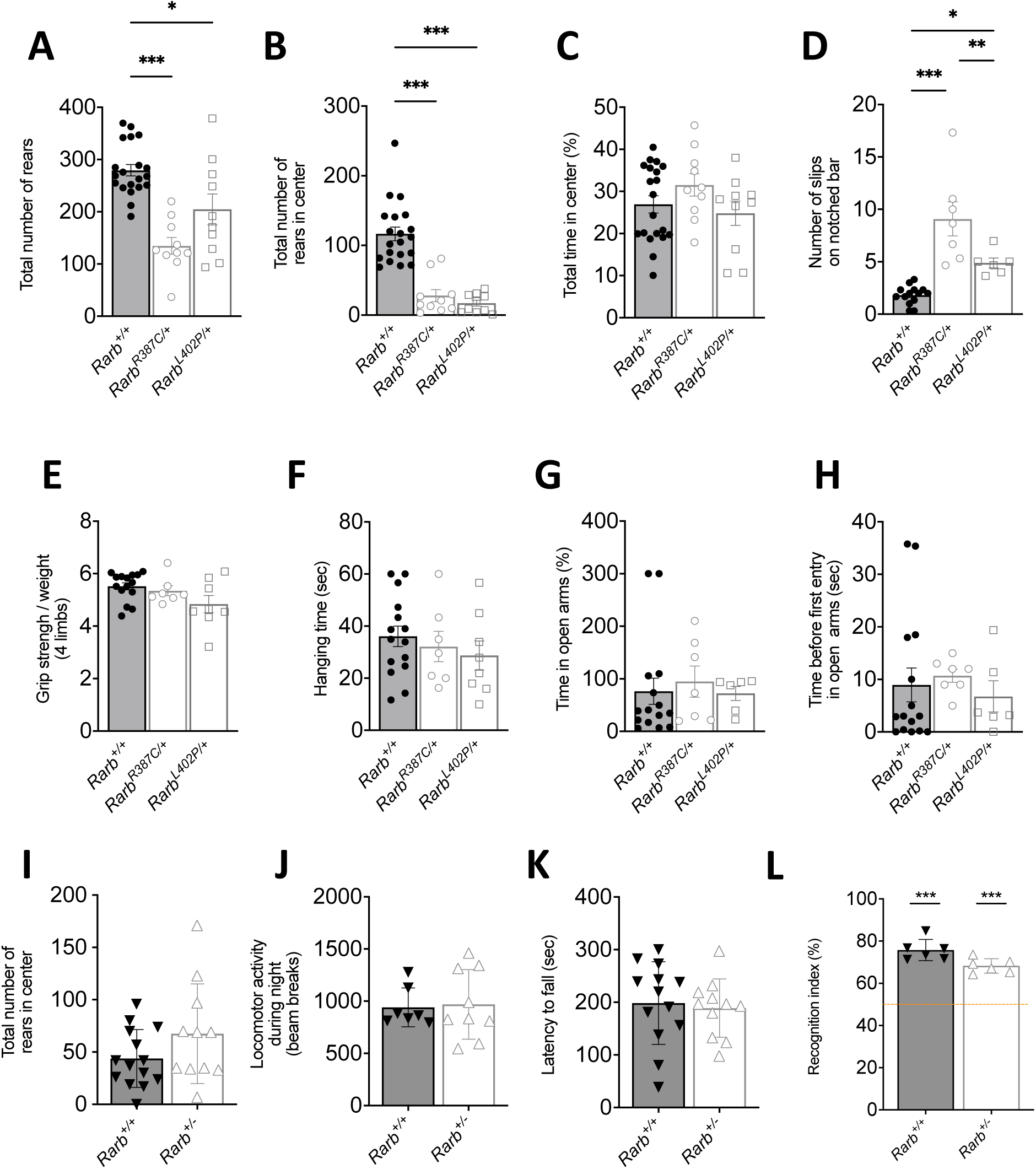
Behavioral performance of RARB-RD mice. (A) Quantification of the total number of rears during 30 minutes of the open field test. (B) Quantification of the total number of rears in the center of the open field arena for 30 minutes. (C) Evaluation of the percentage of time spent in the center of arena for 30 minutes (note that no difference was observed during the first 5 minutes of the test). (D) Motor coordination was assessed as number of slips on a 1 meter notched bar and it was represented as mean of 3 trials. (E) Muscular strength measurements were assessed as mean of grip strength corrected for body weight or (F) by measuring hanging time of mice on a inverted grid. (G) Anxiety was evaluated in elevated plus maze by measuring the time spent in anxiogenic open arms and (H) the elapsed time before the first entry in anxiogenic open arms. (I) Quantification of the total number of rears during 30 minutes in the center of open field arena for *Rarb*^+/-^ mice. (J) Spontaneous locomotor activity was measured in actimetric cages during 12 hours of the dark cycle for *Rarb*^+/-^ mice. (K) *Rarb*^+/-^ mice motor coordination was evaluated in accelerated rotarod as mean latency to fall from the rotating cylinder. (L) *Rarb*^+/-^ mice short-term declarative memory was assessed with novel object recognition test. Recognition index was then calculated and plotted, and orange line indicates at chance performance level. Statistical differences were calculated using Bonferroni multiple comparisons as post-hoc follow-up of ANOVA analyses or Mann-Whitney test and indicate: *, p < 0.05; * *, p < 0.01; * **, p < 0.001. Error bars represent SEM.

**Figure S4.**
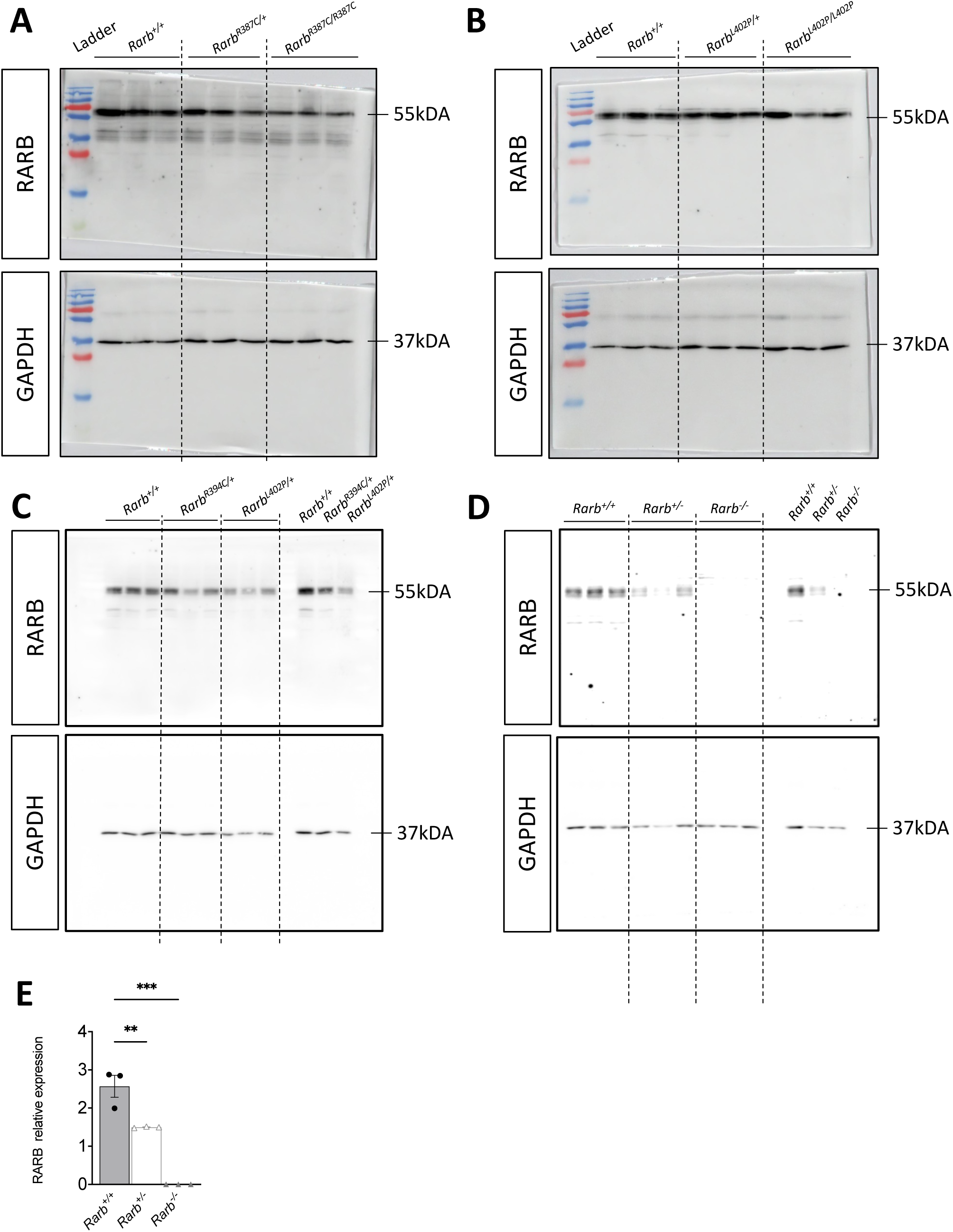
Western blot detection of RARB expression. (A) Western blot detection of RARB and GAPDH (full gels) in the striatum samples from newborn mice heterozygous and homozygous for *Rarb R387C* (left pannels) and (B) *Rarb L402P* (right panels) mutant alleles. (C) Western blot detection of RARB and GAPDH (full gels) in the striatum samples from heterozygous *Rarb R387C* and *Rarb L402P* adult mice. (D) Western blot detection of RARB and GAPDH (full gels) in the striatum samples from *Rarb^+/+^* (WT), *Rarb^+/-^ and Rarb^-/-^* mice and (E) its quantification. RARB expression was normalized by GAPDH expression to control for input quantity. Statistical differences indicate: *, p < 0.05; * *, p < 0.01; * **, p < 0.001. Error bars represent SEM.

**Figure S5.**
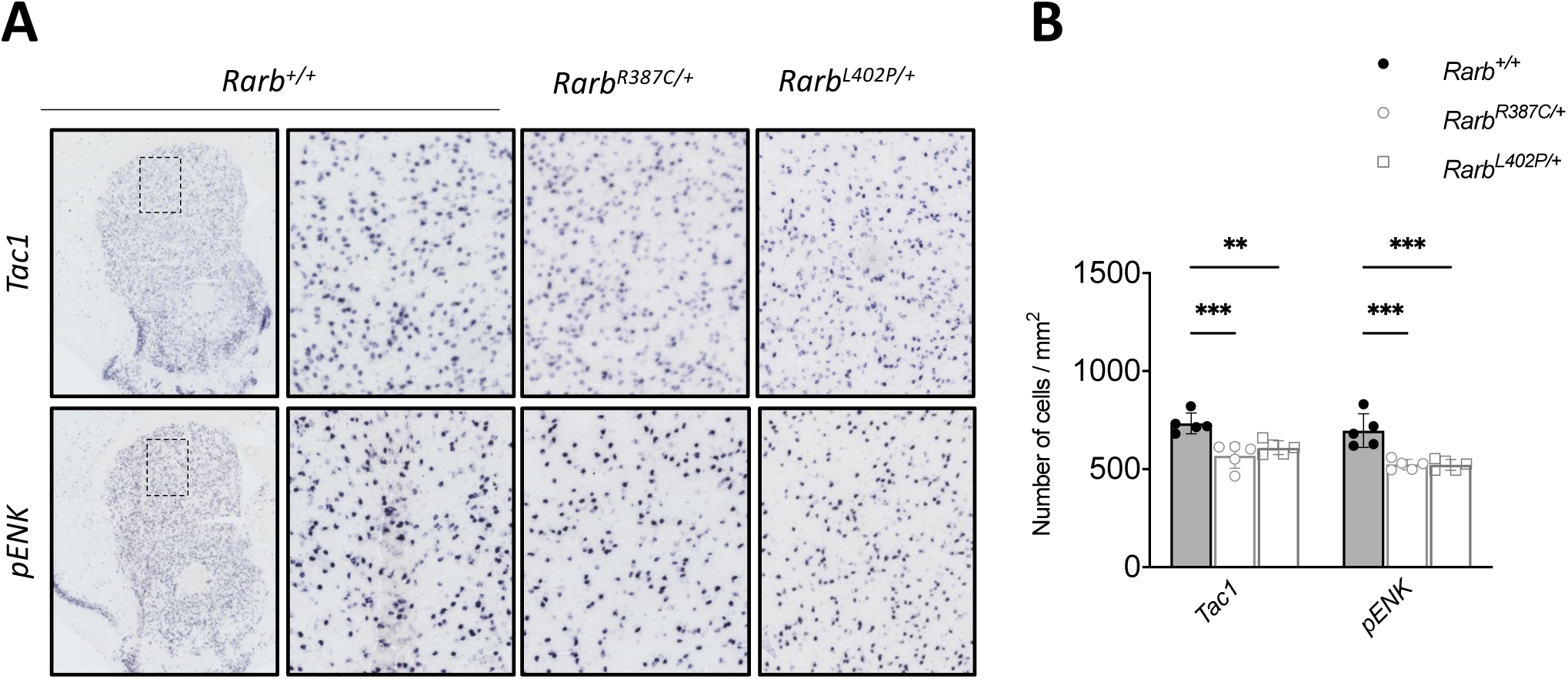
Analyses of markers of MSN subtypes. (A) Examples of *in situ* hybridization identification of snMSNs (*Tac1+*) or spMSNs (*pENK+*). Selected regions of dorsolateral striatum (see rectangles) were enlarged in adjacent panels and correspond to the areas used for scoring number of cells positive for *Tac1* and *pENK* shown in (B). Statistical differences were calculated and indicate : *, p < 0.05; **, p < 0.01; ***, p < 0.001. Error bars represent SEM.

**Fig. S6.**
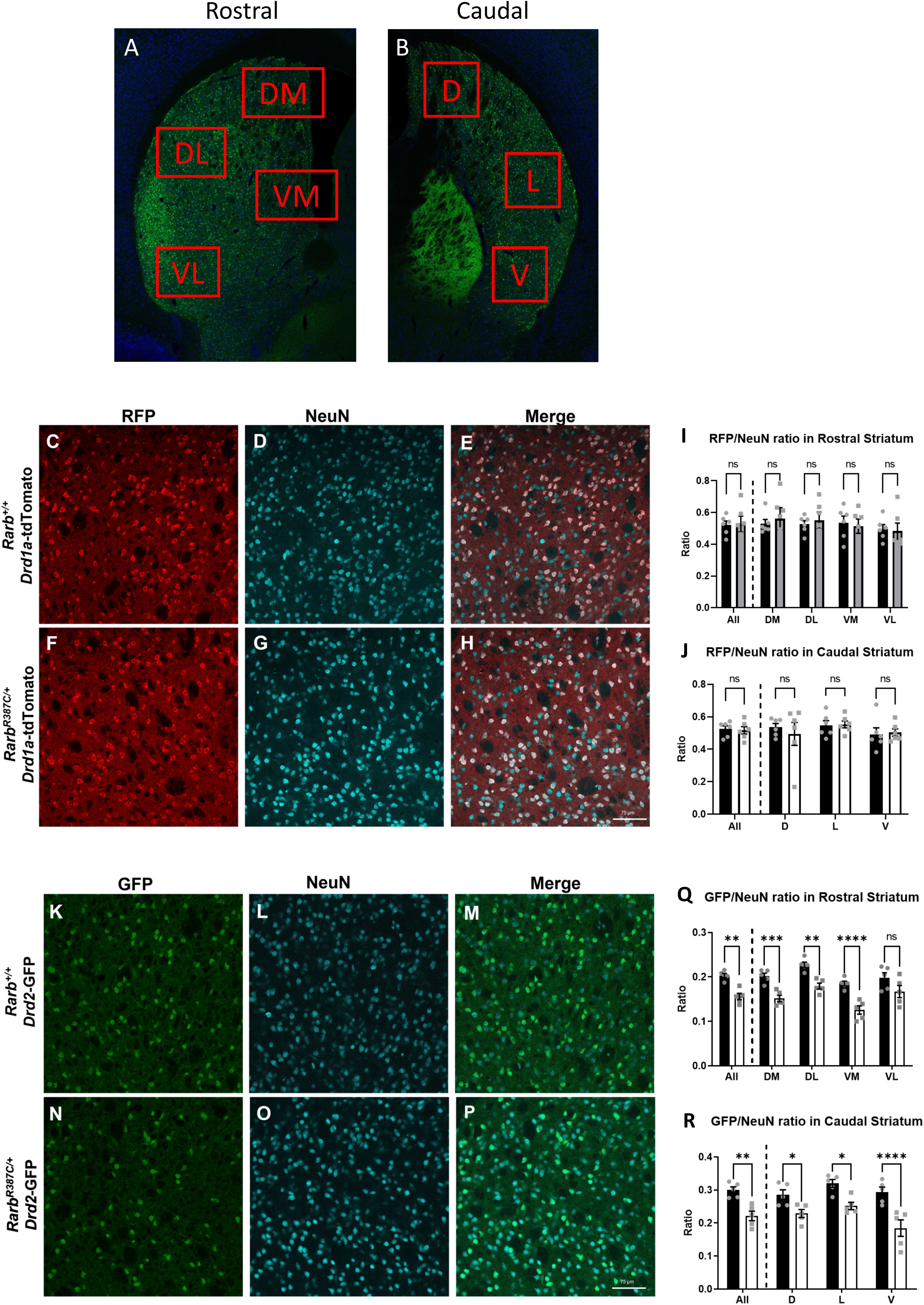
Quantification of the number of neurons expressing the *Drd1a-*tdTomato or *Drd2-*GFP reporter transgenes in the striatum of *Rarb^+/+^* and *Rarb^R387C/+^* mice. (A,B) For cell quantification, sampling boxes (250 μm x 250 μm) were placed in four rostral areas (DM: dorsomedial; DL: dorsolateral; VM: ventromedial; VL: ventrolateral) and three caudal regions (D: dorsal; L: lateral; V: ventral) of the dorsal striatum. (C-H) Representative images of the expression of RFP and NeuN markers, as well as their colocalization *(merge) in Rarb^+/+^* (C-E) and *Rarb^R387C/+^* (F-H) in 2-month-old mice. (I,J) Ratio of the number of cells expressing RFP over NeuN-expressing cells in the rostral (I) and caudal (J) striatum for each genotype. (K-P) Representative images of the expression of GFP and NeuN markers, as well as their colocalization *(merge)* in *Rarb^+/+^* (K-M) and *Rarb^R394C/+^* (N-P) in 2-month-old mice. (Q,R) Ratio of the number of cells expressing GFP over NeuN-expressing cells in the rostral (Q) and caudal (R) striatum for each genotype. Data from 6 *Rarb^+/+^* and 6 *Rarb^R387C/+^* were used for the *Drd1a*-tdTomato experiments and 5 *Rarb^+/+^* and 5 *Rarb^R387C/+^* for the *Drd2*-GFP experiments. Each datapoint represents the averaged ratio or number of cells from corresponding ROIs of 3 consecutive striatum slices for each animal. Genotypes were compared using Unpaired t-tests followed by Welch’s correction. *, p < 0.05; **, p < 0.01; ***, p < 0.001; ****, p < 0.0001 compared to WT littermates. ns: not significant. Scale bar: 75 µm.

**Fig. S7.**
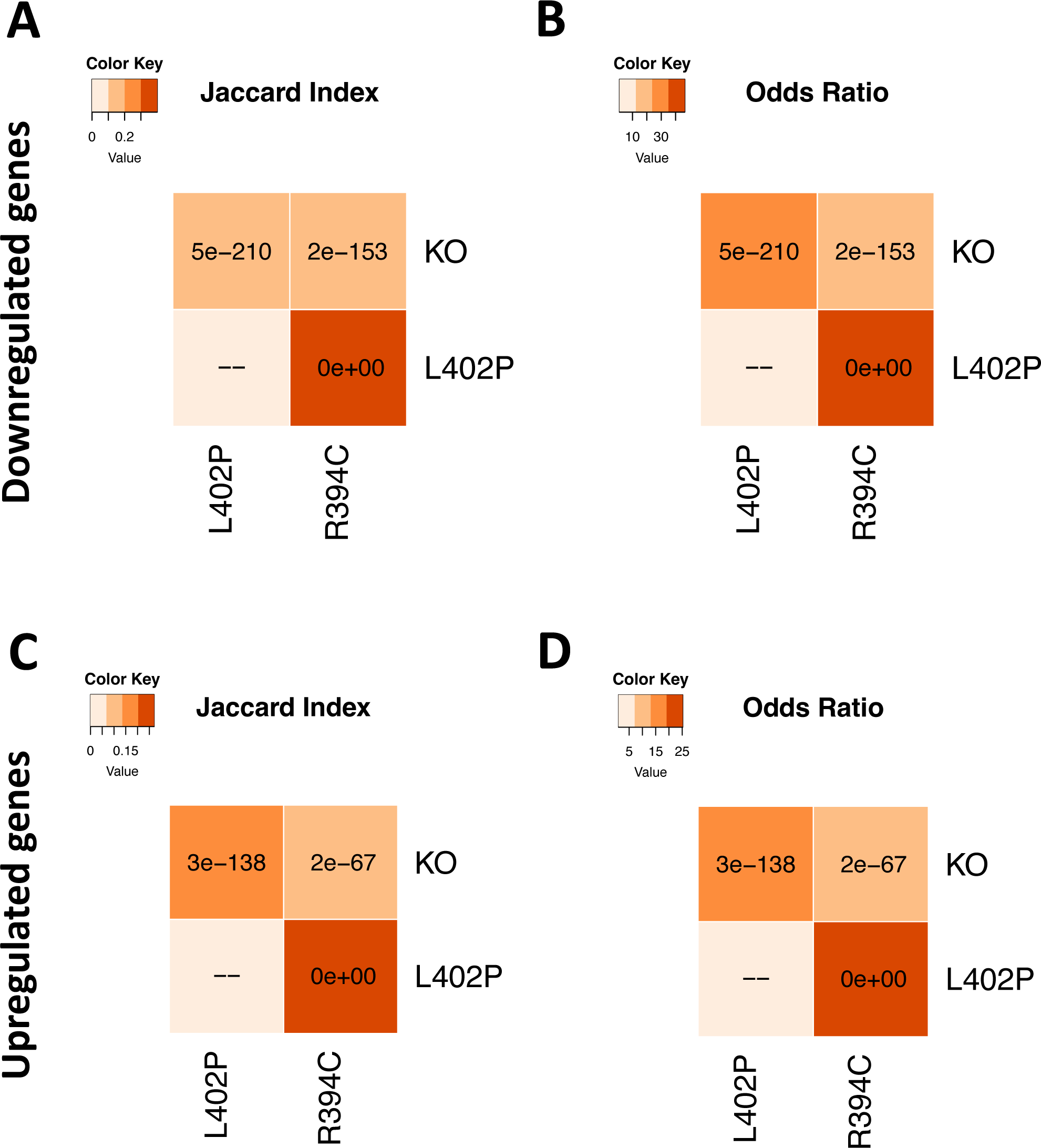
Genes overlap between *Rarb* mutant mice. The *GeneOverlap* tool was used to assess whether the overlap between genes that are downregulated (A, B) or upregulated (C, D) in *Rarb*^R387C/+^, *Rarb*^L402P/+^ or *Rarb*^-/-^ mice is statistically significant. *GeneOverlap* applies Fisher’s exact test to estimate statistical significance and calculate the odds ratio. P-values are indicated within the relevant elements of the matrices, while Jaccard indices (A, C) and odds ratios (B, D) are represented by the color filling each matrix element.

**Figure S8.**
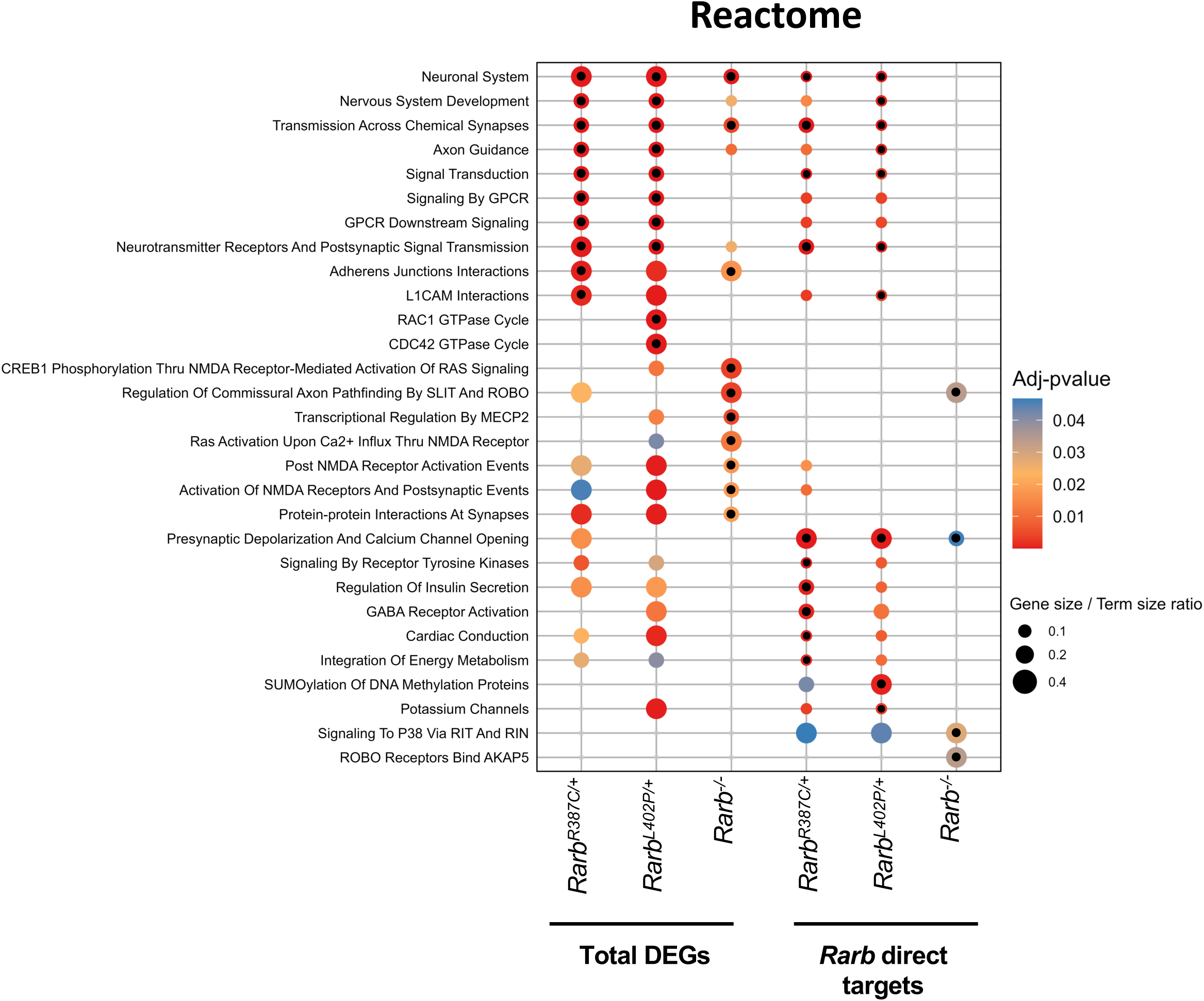
Pathway and gene set enrichment in the transcriptomes of RARB-RD and *Rarb^-/-^*mice. (A) Reactome analysis of all DEGs or of downregulated genes encoding RARB targets in *Rarb^R387C/+^*, *Rarb^L402P/+^* or *Rarb^-/-^* mice, indicating significant enrichment of similar functional terms in RARB-RD mice. Adjusted p-values (p < 0.05) are indicated by the color code and gene/term ratio by the size of the dots. The top 10 significant pathways for each line are indicated by the black dots within the colored ones.

